## Supplementary material for "Optochemical control over mRNA translation by photocaged phosphorodiamidate morpholino oligonucleotides *in vivo*": Tarbashevich et al_Supplemental_info

---

[1]Institute of Cell Biology, Center for Molecular Biology of Inflammation (ZMBE), Muenster, Germany.

[2]Max Planck Institute for Molecular Biomedicine, Münster, Germany.

[3]School of Applied and Interdisciplinary Sciences, Indian Association for the Cultivation of Science, Jadavpur, Kolkata 700 032, India

*

**-Supplemental Figures 1-26 and Figure Legends**

**-Supplemental Tables 1-4**

**-Supplemental Schemes 1-2**

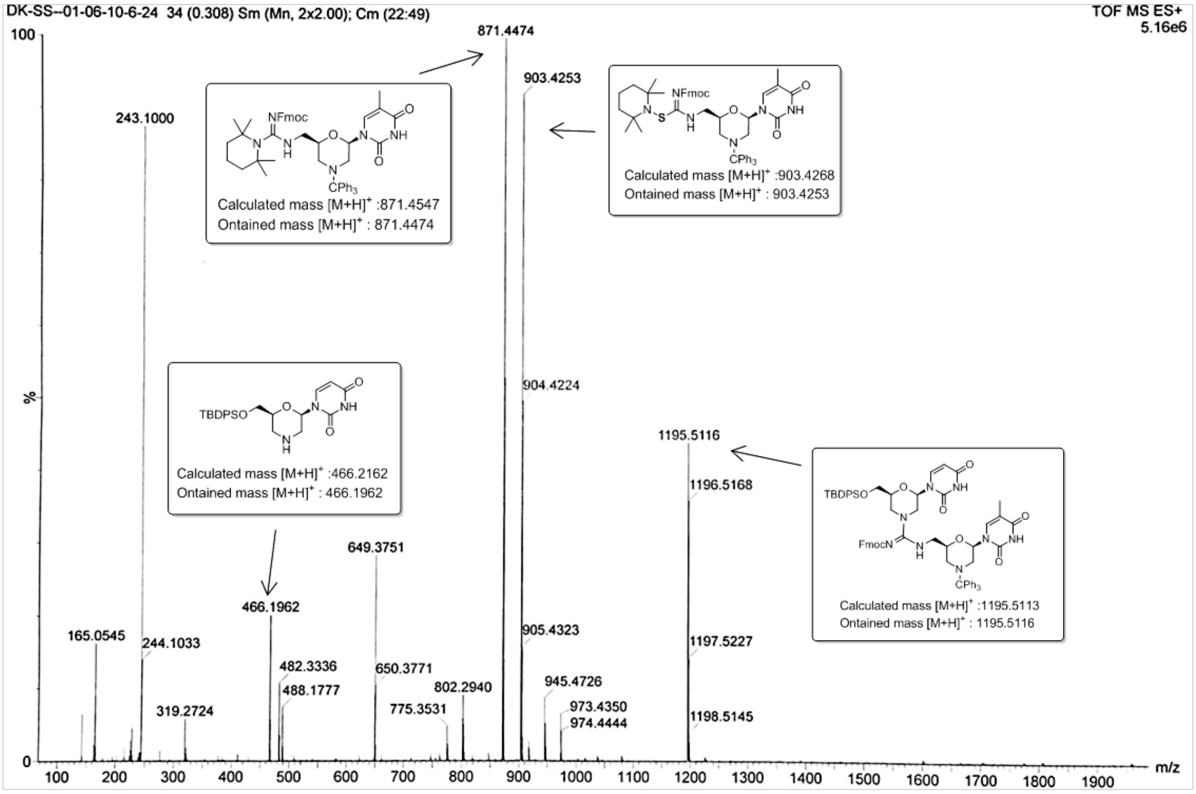


**Figure S1.** Mass spectra of crude reaction mixture by $I_2$/TMP

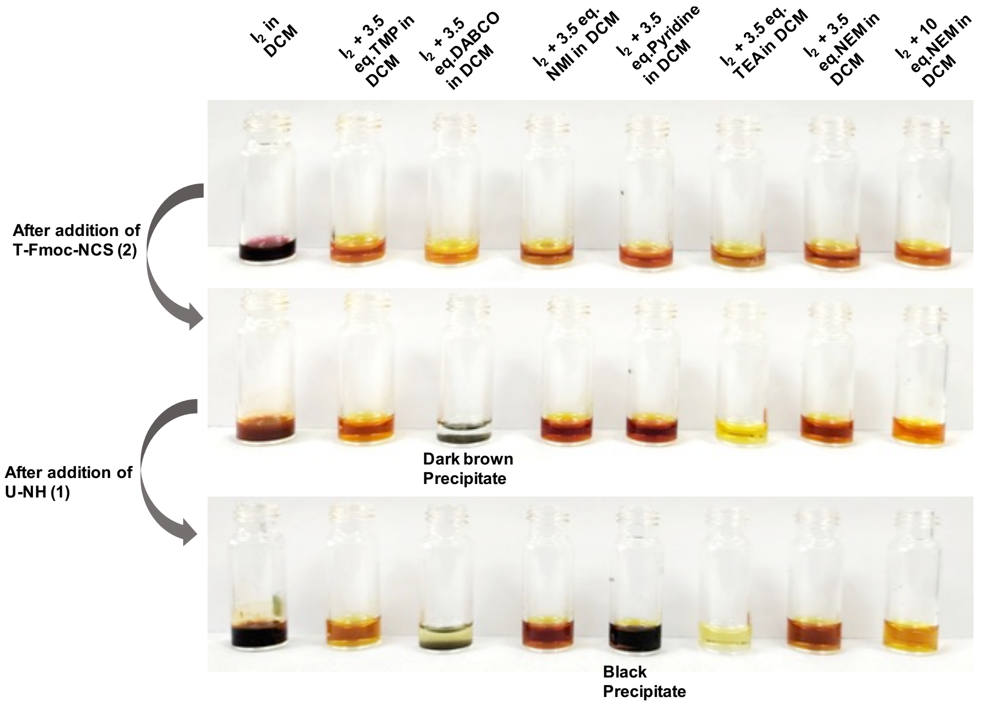


**Figure S2.** Images of the reaction mixtures in presence of different bases

#### Scheme S1: (A) Synthesis of photo caged thymidine monomer. (B) Solid phase synthesis of GMO.

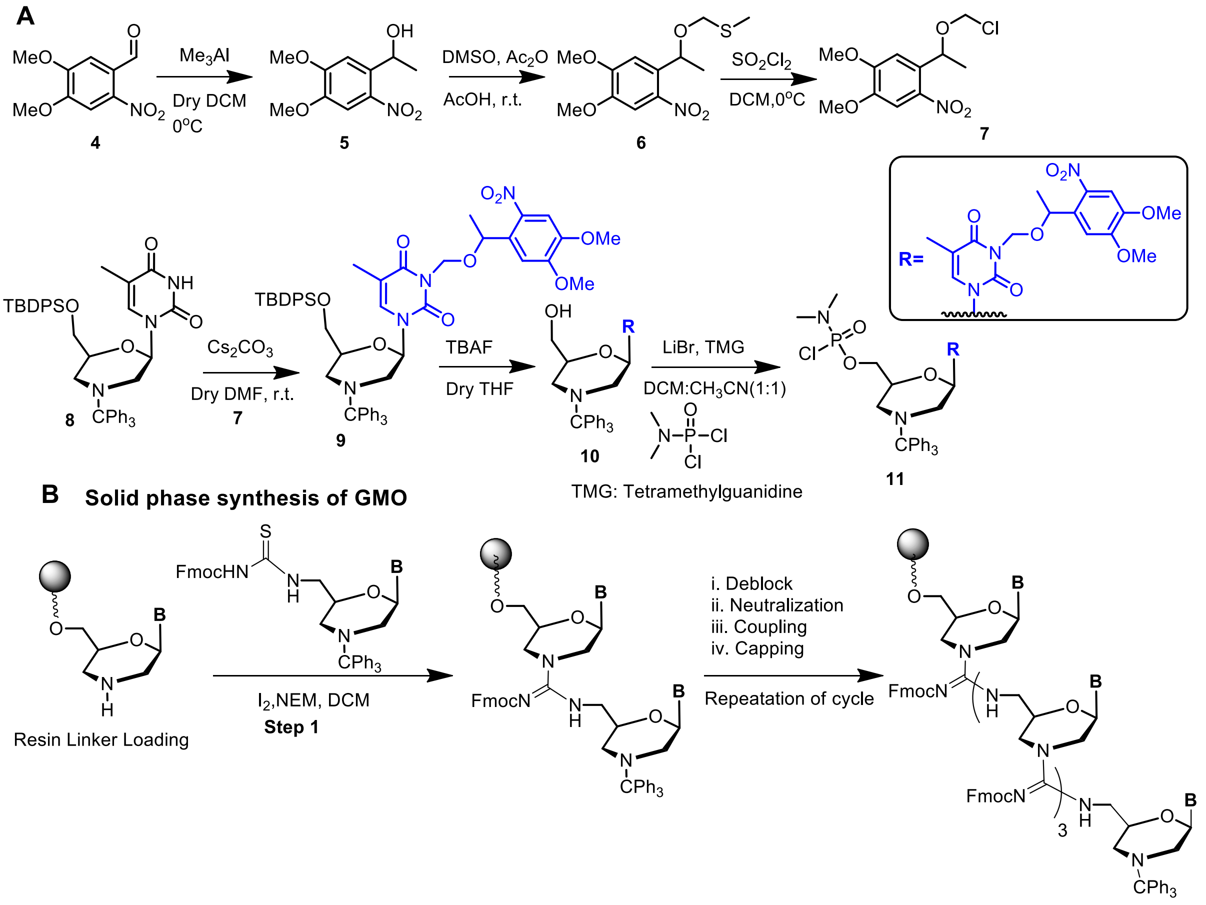

Reagents and conditions: (i) Deblock: CYPMSA (2% salt of 3-cyno pyridine and methane sulphonic acid) in 20% MeOH-DCM (ii) Neutralization: 20% DIPEA-NMP (iii) Coupling: T-Fmoc-NCS, $I_2$, NEM, DCM solvent (iv) 20%-DIPEA-NMP and 20% $Ac_2O$-NMP in 1:1 mixture

**Manual synthesis of GMO on solid support**

In order to synthesis the photo-caged GMO-PMO, the GMO part was synthesized manually using Ramage chemmatrix resin. For the synthesis of guanidinum linkage Fmoc thiourea morpholino monomers were used and coupling was done in presence of $I_2$ and NEM in DCM solvent.

**Scheme S2: (a) PMO synthesis cycle in automated synthesizer and Synthesized PMOs**

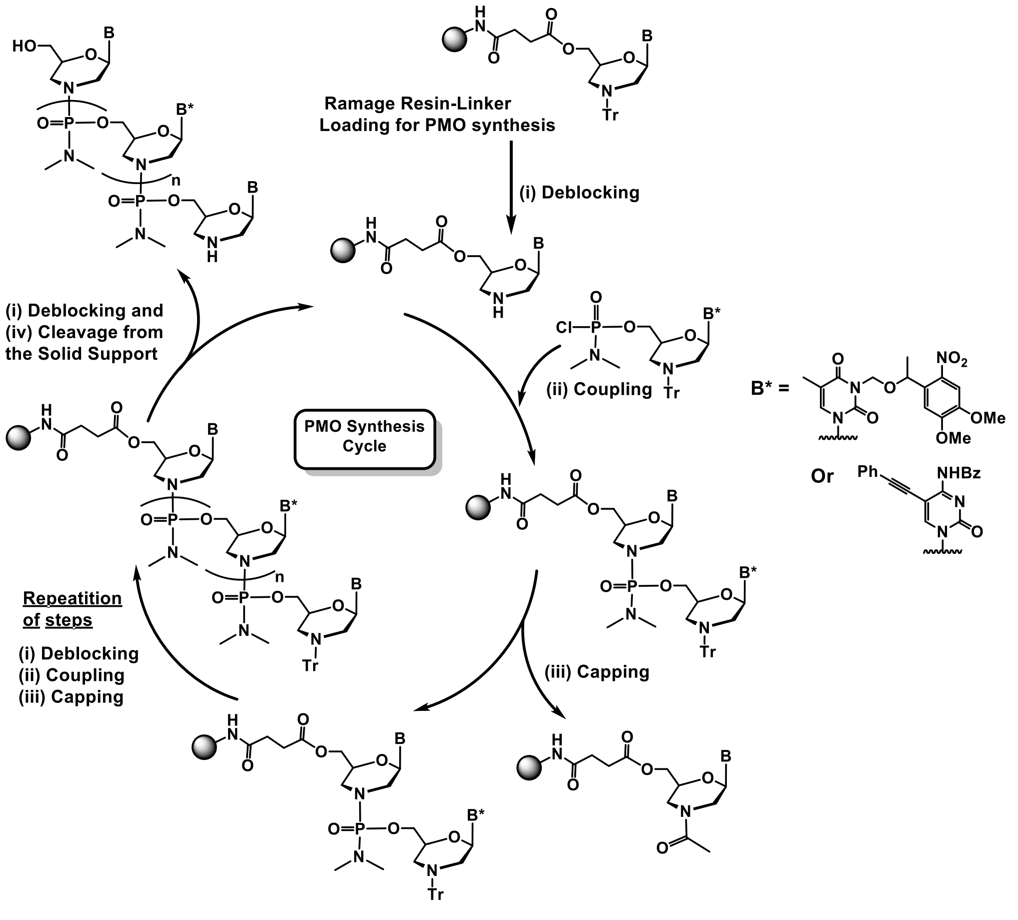


Reagents and conditions: (i) (a) Deblock: CYPMSA (2% salt of 3-cyno pyridine and methane sulphonic acid) in 20% MeOH-DCM (b) Neutralization: 20% DIPEA-NMP (ii) Coupling: Chlorophosphoramidate monomer, ETT, NEM, NMP solvent (iii) Capping: 20%-DIPEA-NMP and 20% $Ac_2O$-NMP in 1:1 mixture (iv) Cleavage from solid support aq. $NH_3$, 55˚C, 16 hr.

#### (b) Synthesis of GMO-PMO chimera in automated synthesizer

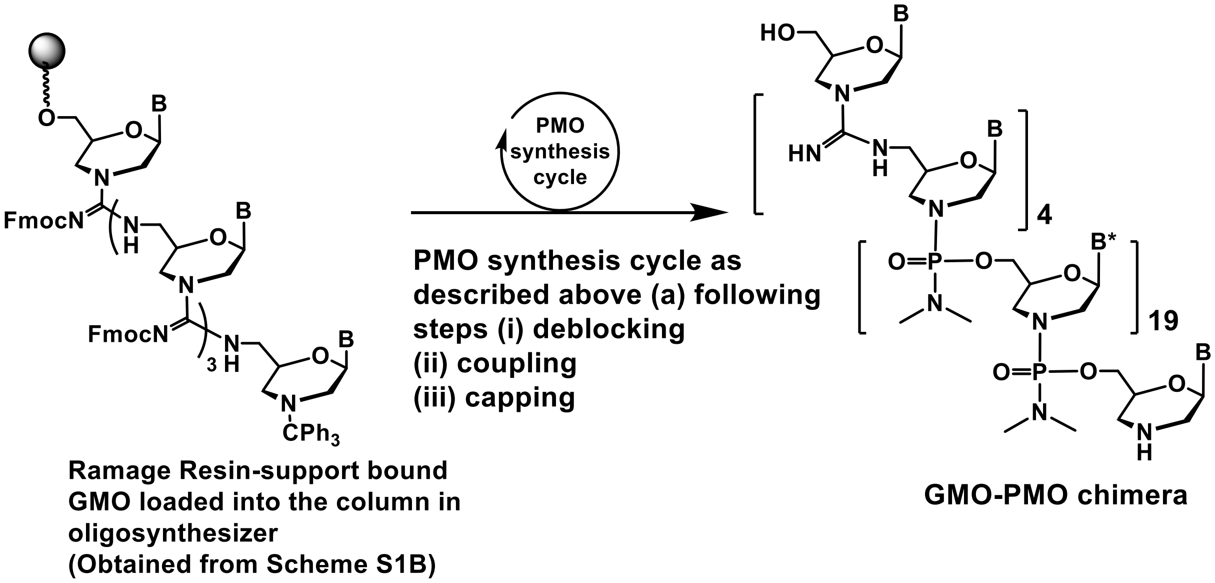


Semi-automated synthesis of GMO-PMO chimera

**PMO synthesis method in automated synthesizer:**

The synthesis column was filled with solid support (Ramage chemmatrix resin) after loading with monomer and connected with oligosynthesizer (H-8, K & A Laboratory, Germany). Loading monomer was attached with solid support via succinic ester linkage which was cleaved by aq. $NH_3$ treatment ([1],[2]).

**Description of the Oligosynthesizer (H-8, K & A Laborgeraete, Germany)**

There are 6 different reagents bottle connected with the machine which are

- I) Washing
- II) CAP A
- III) CAP B
- IV) Deblocking
- V) Activator
- VI) Oxidizer.

Total 12 amidite bottles can be connected at a time. (i) 4 normal positions (A, T, G and C) for normal amidites, (ii) 4 modified positions (Z, O, S and U) for modified amidites, (iii) 4 additional positions (1, 2, 3 and 4) for very small scale synthesis. For the synthesis of PMO normal A, T, G and C positions were accessed. There was no need for oxidizer. An additional DCM (Dichloromethane) wash was required prior to each deblocking which was performed using oxidizer bottle. U (one of the four available modified amidite positions) was used for delivering neutralizing reagent. Each automated synthesis cycle is composed of four steps excluding washing.

I) **Washing:**

**Step 1-**Gas flow for 2s through the column from the bottom to the waste.

**Step 2-**200 μL DCM was purged with 2s delay through the column from the bottom to the waste.

**Step 3**-Repeat of **Step 1** and **Step 2** for another two times.

**Step 4**-Gas flow for 0.5s through the valve block to the waste with 1s delay.

II) **Deblocking:**

**Step 1-**Gas flow for 4s through the column from the bottom to the trityl monitor with 1s delay.

**Step 2**-200 μL of deblocking cocktail CYPMSA (2% 3-cyanopyridine and methanesulfonic acid salt) was purged with 30s delay through the column from the bottom to the trityl monitor.

**Step 3**-Repeat of **Step 2** for another 2 times.

**Step 4**-200 μL DCM was purged with 1s delay through the column from the bottom to the waste.

**Step 5**-Gas flow for 2s through the column from the bottom to the waste repeat the step for two times.

III) **Neutralization:**

**Step 1**-100 μL of neutralizing solution (20 % DIPEA-NMP) was purged with 30 s delay.

**Step 2-**Repeat of Step 1 for another time.

**Step 3-** Gas flow for 0.5s through the valve block to the waste with 1s delay.

IV) **Coupling:**

**Step 1**-Gas flow for 3s through the column from the bottom to the waste with 1s delay.

**Step 2**-Gas flow for 0.5s through the valve block to the waste with 1s delay.

**Step 3**-50 μL of active monomer (0.2 M) and 20 μL of ETT (5-Ethylthio-1H-Tetrazole) (0.3 M) /NEM (N-ethyle morpholino) (0.5M) were purged with 9 min delay through the column from the bottom to the waste.

**Step 4**-Repeat of Step 3 for another 2 times

**Step 5**-Gas flow for 10 s through the column from the bottom to the waste.

**Step 6**-100 μL DCM was purged through the valve block to the waste.

**Step 7**-100 μL $CH_3CN$ was purged through the column from the bottom to the waste.

**Step 8**-Repeat of **Step 1** and **Step 2** for single time.

V) **Capping:** This was performed using '1ucap' subroutine.

**Step 1**-50 μL of **CAP A** (20% $Ac_2O$-NMP) and 50 μL of **CAP B** (20% DIPEA-NMP) were purged with 30 s delay through the column from the bottom to the waste.

**Step 2**-Repeat of **Step 1** for two times

**Step 3**- Gas flow for 3s through the column from the bottom to the waste.

**Step 4**-100 μL $CH_2Cl_2$ was purged through the valve block to the waste.

**Step 5**-100 μL $CH_3CN$ was purged through the column from the bottom to the waste.

**Step 6**- Gas flow for 3s through the valve block to the waste with 1s delay.

After synthesis solid support was washed with DCM and dried by purging with argon and deprotected from solid support using 30% aqueous ammonia at 55°C for 16h.

**Table S1**: List of GMO-PMOs and PMOs synthesized*

| | |
|---|---|
| GMOPMO-1 | 5′-T^T^T^T^TTTTTT-3′ |
| GMOPMO-2 | 5′-T^A^T^A^ATTTTT-3′ |
| cPMO 1 | 5′-TATAAATTGTAACTGAGGTAAGAGG-3′ |
| PMO1 | 5′-TATAAATTGTAACTGAGGTAAGAGG-3′ |
| cPMO2 | 5′-T^A^T^A^AATTGTAACTGAGGTAAGAGG-3' |
| PMO 2 | 5′-T^A^T^A^AATTGTAACTGAGGTAAGAGG-3′ |
| PMO 3 | 5′-T^A^T^A^AATTGTAACTGAGGTAAGAGG-3′ |
| tbMO | 5'-CCTCTTACCTCAGTTACAATTTATA-3' |
| PMO1-Bodipy | 5′-TATAAATTGTAACTGAGGTAAGAGG-Bodipy-3′ |
| PMO2-Bodipy | 5′-T^A^T^A^AATTGTAACTGAGGTAAGAGG-Bodipy-3′ |
| PMO3-Bodipy | 5′-T^A^T^A^AATTGTAACTGAGGTAAGAGG-Bodipy-3′ |

*HPLC chromatograms (Table S2) and MALDI TOF mass (Table S3) (*vide infra* page 37 and 44, respectively)

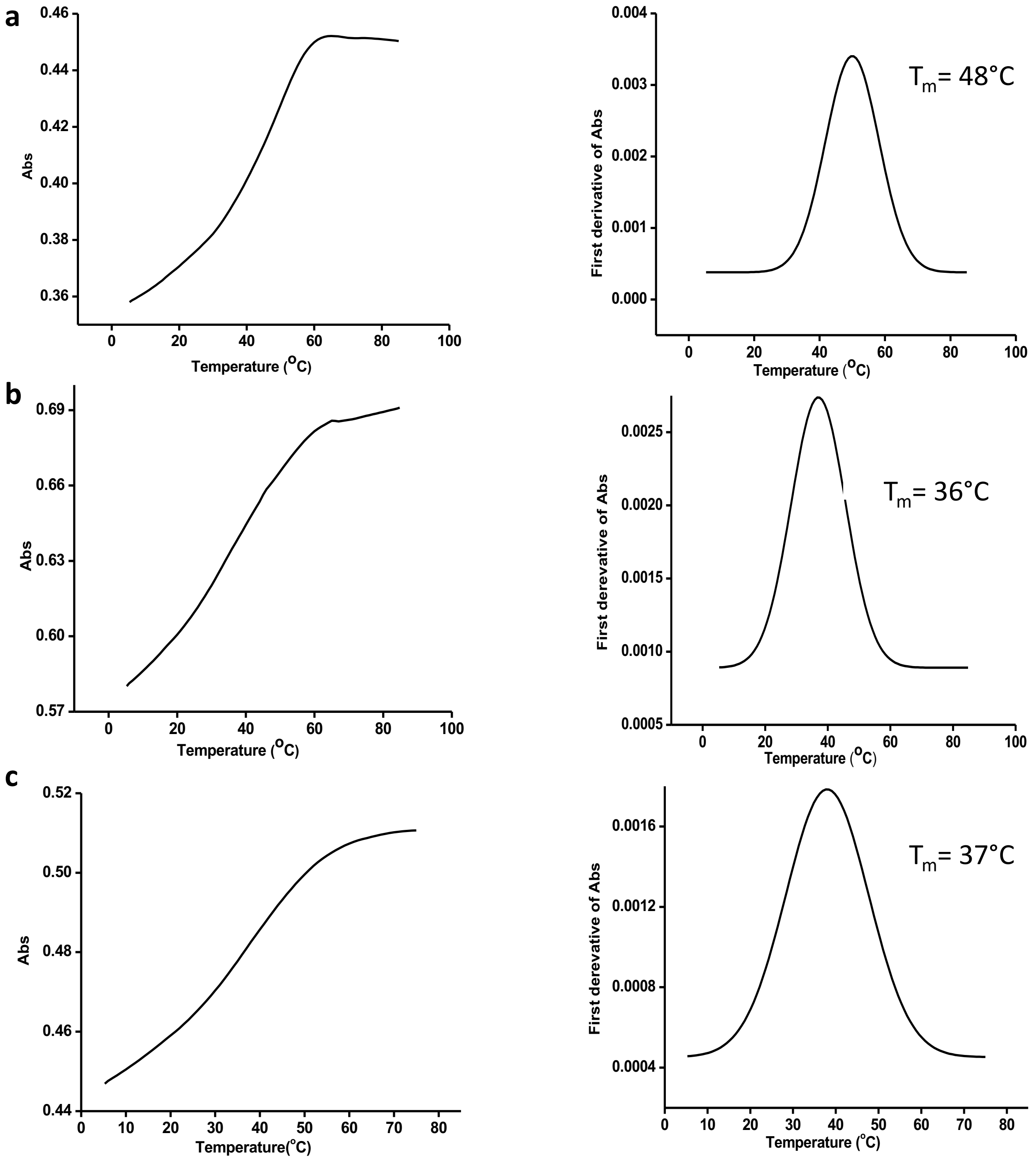


**Figure S3**. Thermal melting curves and its first derivative plot of (**a**) tbMO-RNA (**b**) tbMO-PMO 1 and (**c**) tbMO-PMO3

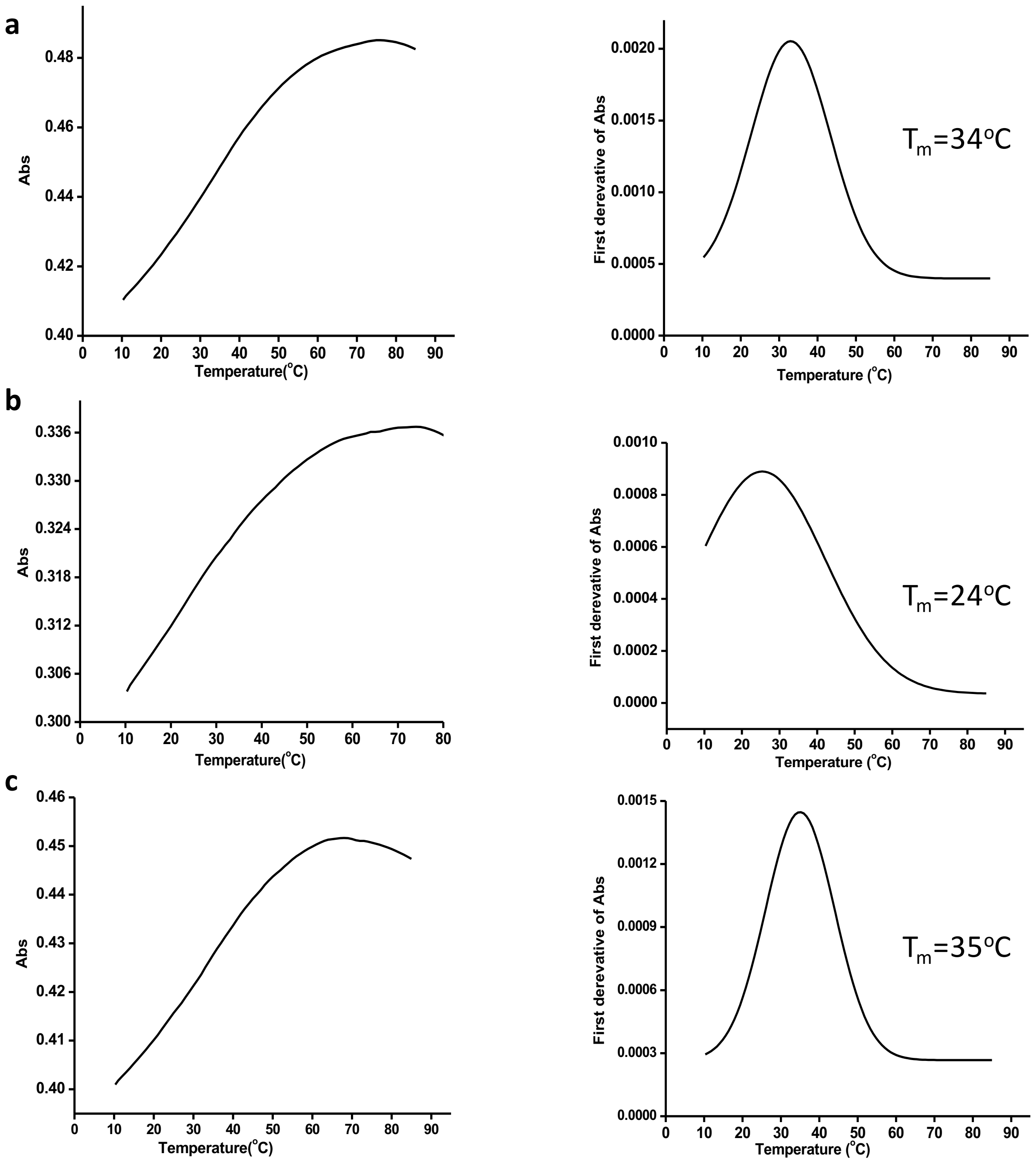


**Figure S4**. Thermal melting curves and its first derivative plot of (**a**) tbMO-PMO2, (**b**) tbMO-cPMO2 (before irradiation) and (**c**) tbMO-PMO2 (after light irradiation)

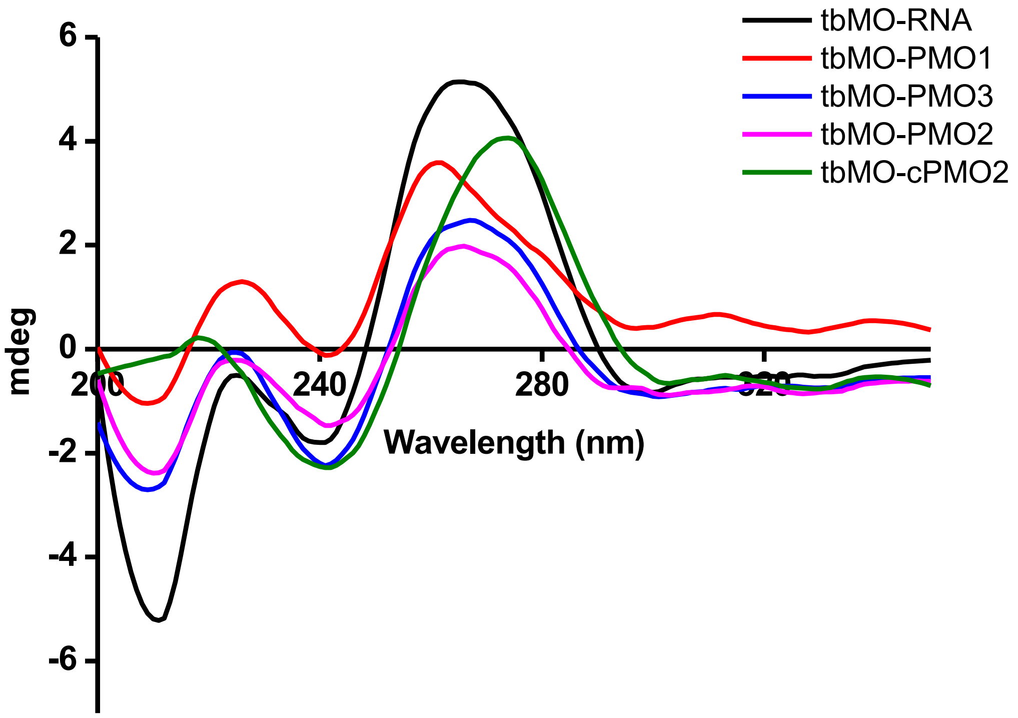


**Figure S5**. CD spectrum of duplexes in 40mM phosphate buffer at 10°C. The duplex samples were obtained by mixing both strands of interest in a 1:1 ratio at 2µm concentration.

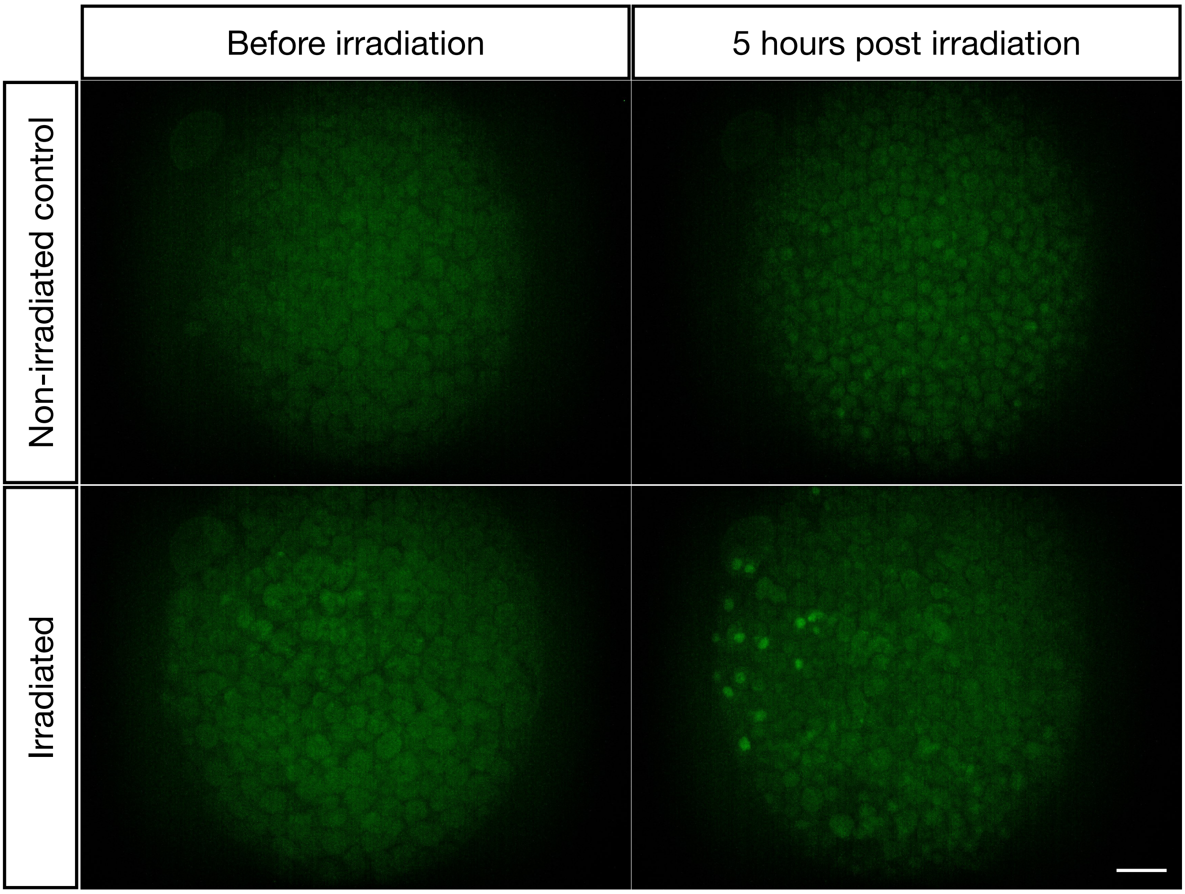


**Figure S6.** Local light-induced expression of GFP as visualized at low magnification. Zebrafish embryos were injected with the mRNA encoding for the nuclear GFP, cPMO2, as well as with the standard translation-blocking morpholino. The embryos were allowed to develop until 4 hours post fertilization (hpf). Some of the embryos were kept in the dark (upper panels), such that the translation of the mRNA was inhibited by the translation-blocking morpholino. In this case, no expression of nuclear GFP protein was detected. Sibling embryos were subjected to local irradiation with the 405 nm laser (lower panels), resulting in the photolysis of PMO2's light-sensitive moieties. The sequestration of the translation-blocking morpholino allowed the translation of the nuclear GFP, which was visualized 5 hours post-irradiation (green spots). Scale bar 100 μm.

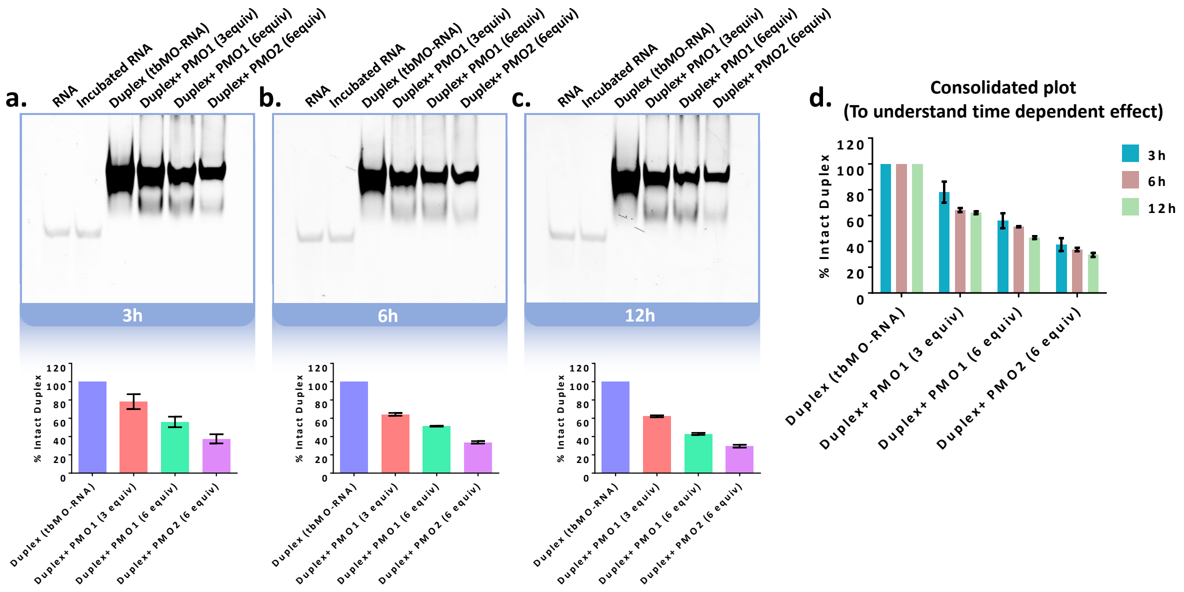


**Figure S7.** Time dependent electrophoretic mobility shift assay for strand displacement. PAGE image (Ethidium Bromide) of tbMO-RNA duplex with PMO1 and PMO2 following (**a**) 3h, (**b**) 6h, (**c**) 12h of incubation at 30 °C with their respective bar graph. Duplex concentration was 50 μM (0.5 nmol in 10 μL buffer). For all the cases, the duplex and RNA were incubated and annealed in PBS 1X buffer (137 mM NaCl, 2.7 mM KCl, 10 mM $Na_2HPO_4$, and 1.8 mM $KH_2PO_4$). Lane 1 contains RNA single strand which was loaded instantaneously. Lane 2 contains RNA single strand which was incubated for the same time period as the other duplexes. Lane 3 represents the duplex after incubation whereas Lanes 4-6 contain the duplex with strand displacing PMOs after incubation. **(d)** Time dependent consolidated plot of percentage of the intact duplexes. Data represent mean ± SD, n= 2.

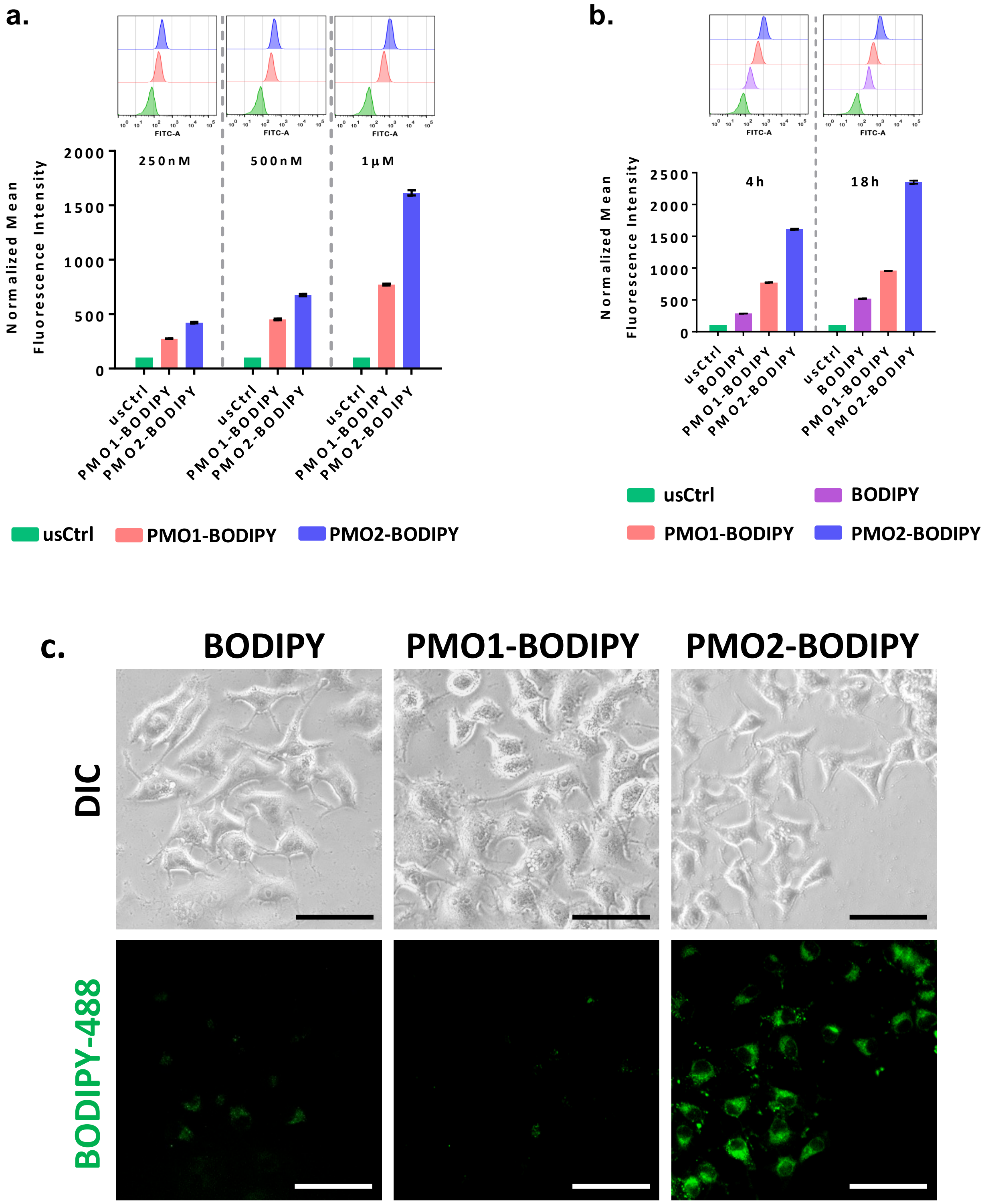


**Figure S8.** Uptake comparisons of Flurophore BODIPY-488, PMO1-BODIPY-488 and PMO2- BODIPY-488 conjugates by flow cytometry in HeLa cell line (**a**) Dose-dependent uptake histograms at 250nM, 500nM and 1μM with their respective bar graph. (**b**) Time-dependent uptake histograms at 1 μM at 4h and 18h with their respective bar graph. Data represent mean ± SD, n= 3. (**c**) Live cell fluorescent microscopy images of BODIPY dye and BODIPY tagged PMOs in HeLa cell line at 1 μM concentration after 24h post treatment. Scale bar 40 μm.

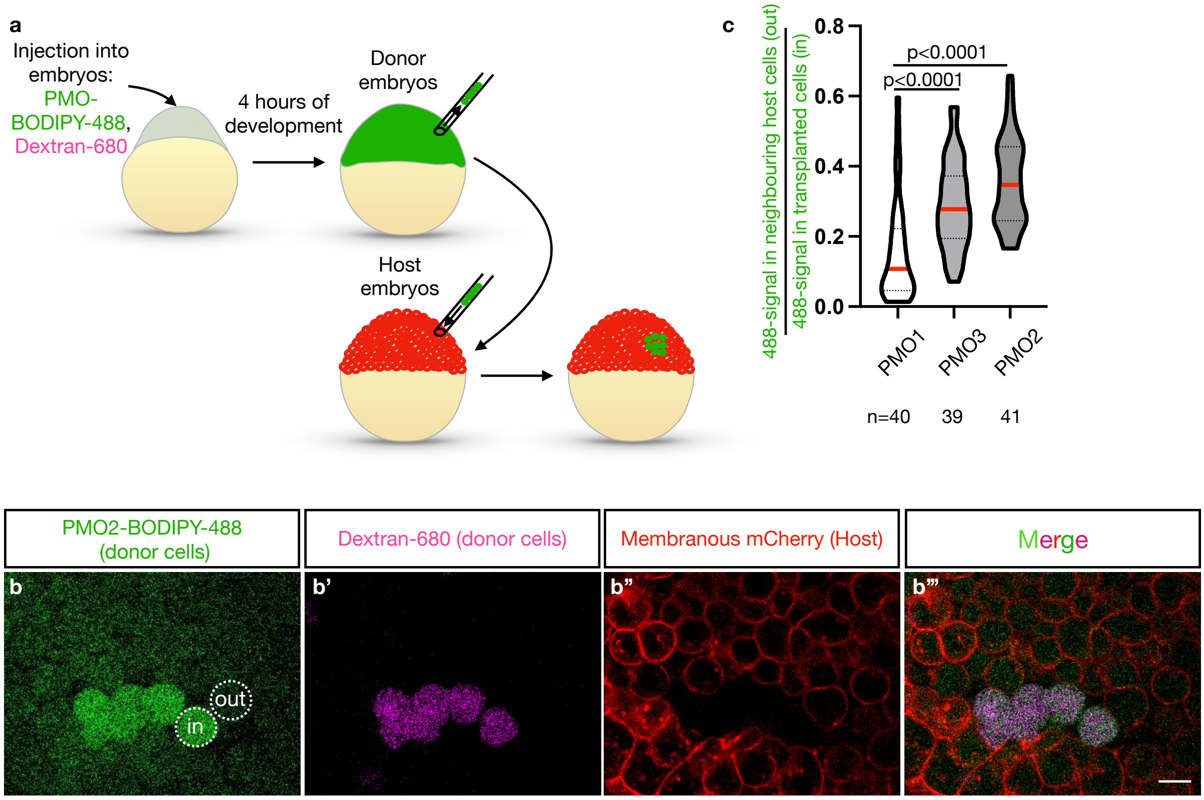


**Figure S9.** PMOs exhibit different diffusion rates through the membrane into neighbouring cells. (**a**) Three different PMOs were labeled with the green BODIPY fluorophore. Donor embryos were injected at 1-cell stage with either of three 10µM fluorescent PMOs (PMO1, PMO2, or PMO3) and 5µM Dextran-680. Cells from these embryos were transplanted into host embryos, where all cell membranes were labeled with mCherry (lower panels). (**b-b'''**) 5µm confocal images were acquired 1 hour after transplantation. (**c**) Quantification of the PMO diffusion was performed by calculating the mean fluorescence ratio of the BODIPY-488 signal within the transplanted cells (marked by Dextran-680) and that in the neighbouring host cells. P-values were determined by ANOVA multiple comparisons test. Dotted circles present an example for the definition of regions from which values were derived. Scale bar 20 µm.

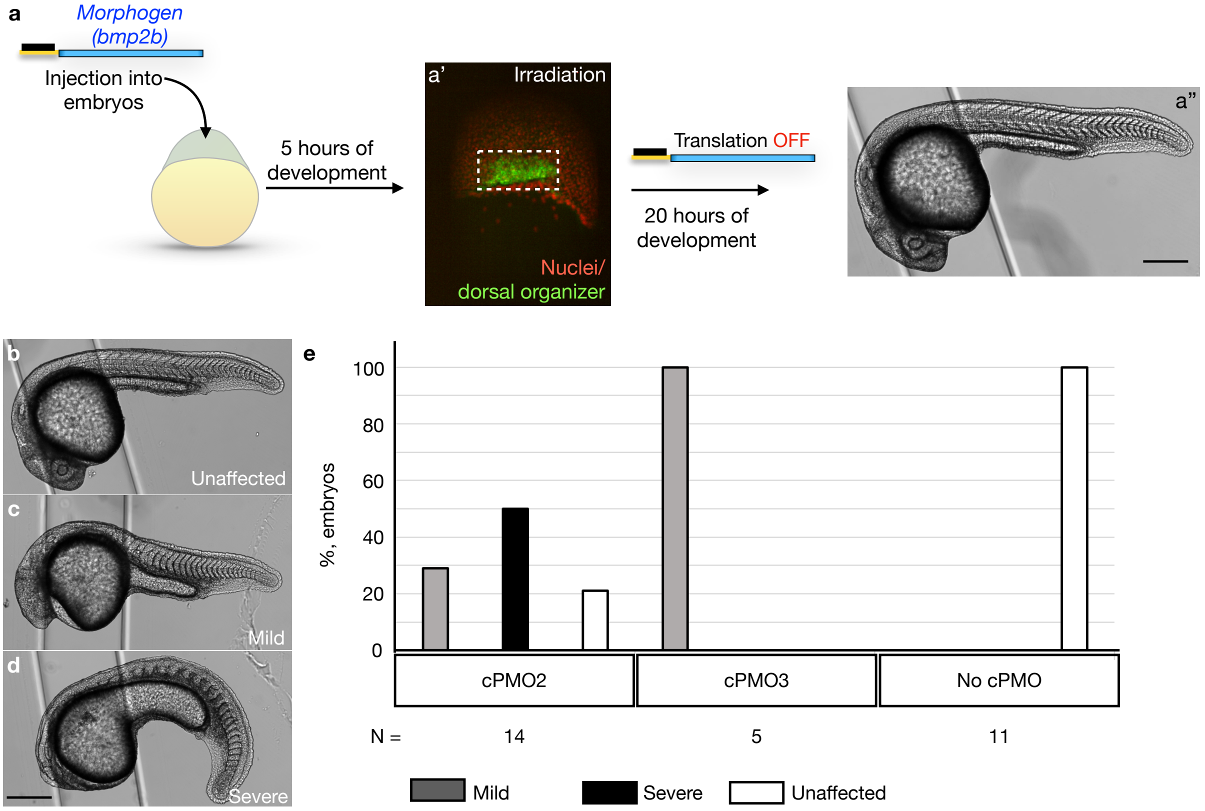


**Figure S10.** Light-induced expression of a Bmp2b morphogen affects zebrafish embryo development. (**a-a''**) Zebrafish embryos were injected with the mRNA encoding for the Bmp2b morphogen (blue) and with the standard translation-blocking morpholino (black). Embryos developed until 5hpf. At this stage, embryos were irradiated locally with the 405nm laser at the dorsal organizer region (green cells in the white rectangle). Bmp2b expression was inhibited by the translation-blocking morpholino and laser irradiation did not affect embryo development. (**b-e**) Embryos were injected with mRNA encoding for a morphogen (Bmp2b) that inhibits the development of dorsoanterior structures, such as the head, cPMO2 or cPMO3, and with the standard translation-blocking morpholino. The embryos were allowed to develop for 5 hours and were then irradiated locally with the 405nm laser at the region of the embryo that induces the development of dorsal structures such as the head, resulting in photolysis of PMO2's light-sensitive moieties (the experiment is equivalent to that presented in **Fig 6a**, lower arrow). This treatment led to the sequestration of the translation-blocking morpholino and the expression of the morphogen that inhibits head development. (**b-d**) Representative images and (**e**) quantification of the severity spectrum of the phenotypes generated by the light-induced Bmp2b mis-expression. Scale bar 400 μm

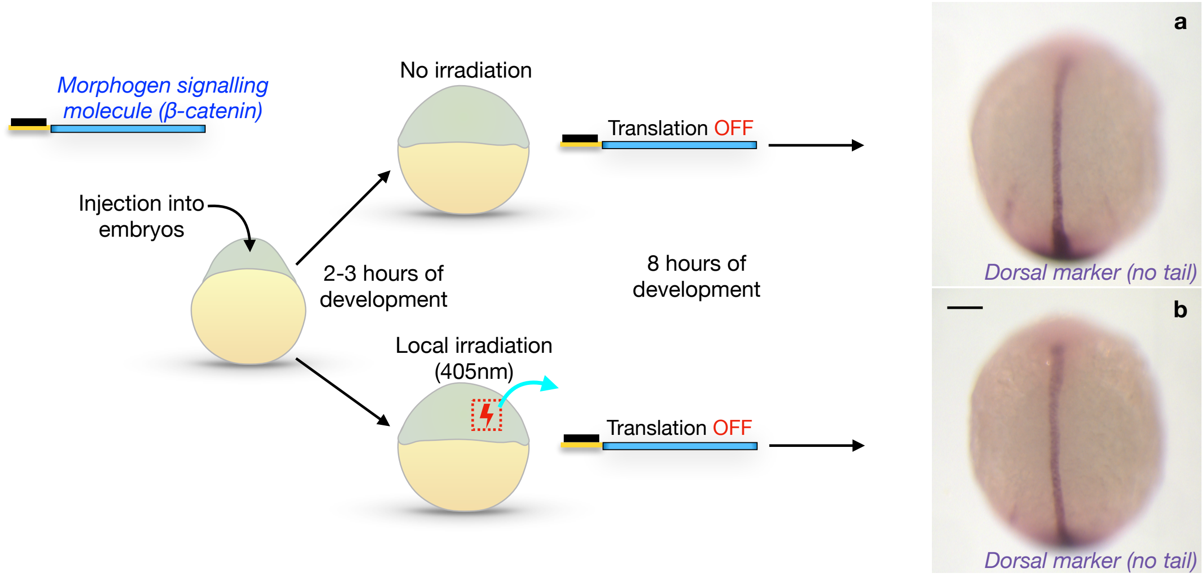


**Figure S11. Light-induced expression of a morphogen downstream signaling factor affects zebrafish embryo development.** (**a-b**) Zebrafish embryos were injected with the mRNA encoding for the ß-Catenin (blue) and with the standard translation-blocking morpholino (black). Embryos developed until 2hpf. At this stage part of embryos were kept in the dark (**a**) and the siblings were irradiated locally with the 405nm laser (**b**). In both cases translation of the mRNA is inhibited by the translation-blocking morpholino, and no morphogen signaling occurs. Dorsal structures were detected by expression of *no tail* (*ntl*) RNA, a dorsal gene marker (violet). Scale bar 150 μm.

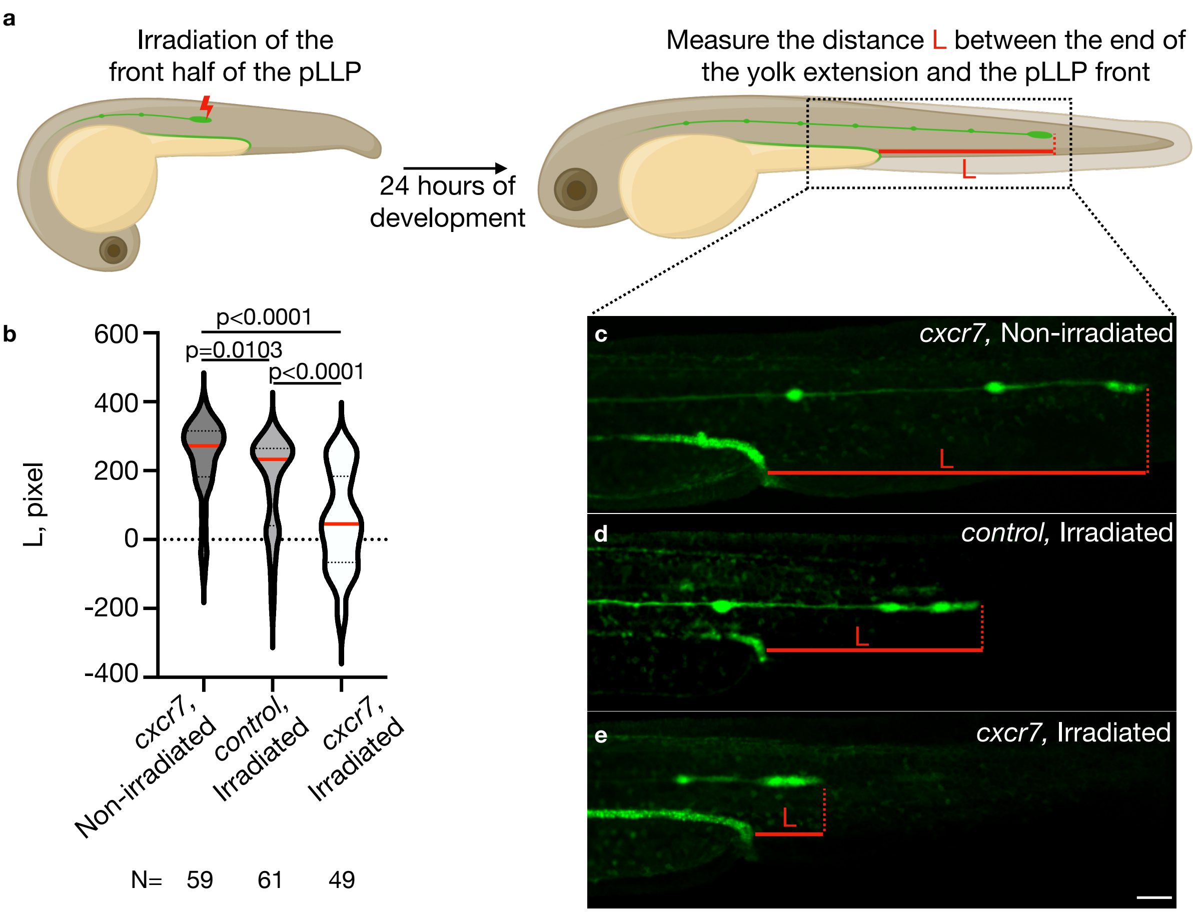


**Figure S12.** Local light-induced expression of the Cxcr7 at the front of the pLLP impairs its migration. (**a**) Embryos were injected with the mRNA encoding for Cxcr7 or control protein, cPMO2, and with the standard translation-blocking morpholino. The embryos were allowed to develop for 24 hours, when part of them was irradiated locally at the front of migrating pLLP with the 405 nm laser, resulting in photolysis of PMO2's light-sensitive moieties. Under these conditions, sequestration of the translation-blocking morpholino allowed the expression of Cxcr7 or control protein to occur. Light-induced expression of the Cxcr7 impairs pLLP migration by inhibiting the generation of the chemokine gradient crucial for the directional movement of the primordium. The pLLP migration phenotype at 48hpf was quantified by measuring the length (L) of the pLLP migration track from the end of the yolk extension to the front of the primordium (**b**). N - number of embryos analyzed. (**c-e**) Representative images of analyzed embryos. Scale bar 200 μm. P-values were determined by ANOVA multiple comparisons test.

**Table S2**: Numbers of embryos analyzed in this work

| Constract | Groups of embryos | Experiment 1 | Experiment 2 | Experiment 3 |
|---|---|---|---|---|
| ***tis-GFPnls*** | cPMO2, irradiated | 18 | 15 | 18 |
| | cPMO2, non-irradiated | 8 | 5 | 5 |
| | cPMO1, irradiated | 10 | 10 | 9 |
| ***tis-kid*** | cPMO2, irradiated | 5 | 10 | 8 |
| | cPMO2, non-irradiated | 3 | 11 | 4 |
| | No cPMO, irradiated | 5 | 10 | 15 |
| | No cPMO, non-irradiated | 3 | 17 | 21 |
| ***tis-bmp2b*** | cPMO2, irradiated | 5 | 5 | 4 |
| | cPMO2, non-irradiated | 8 | 5 | 5 |
| | cPMO3, irradiated | 1 | 2 | 2 |
| | cPMO3, non-irradiated | 1 | 2 | 2 |
| | No cPMO, irradiated | 3 | 5 | 3 |
| ***tis-ctnnb1*** | cPMO2, irradiated | 12 | 15 | 7 |
| | No cPMO, irradiated | 6 | 5 | 5 |
| ***tis-cxcr7*** | cPMO2, irradiated | 19 | 13 | 17 |
| | cPMO2, non-irradiated | 22 | 14 | 23 |
| ***tis-control*** | cPMO2, irradiated | 22 | 25 | 14 |

#### General Synthetic procedures

**Synthesis of GMO dimer (Compound 3) (From Scheme 1)**

Compound 2 (426 mg, 1.2 eq, 0.56 mmol) was dissolved in dry DCM (10 ml) under Ar atmosphere followed by the addition of iodine (220 mg, 2 eq, 0.86 mmol). Then the reaction mixture was cooled in an ice bath and 4-ethylmorpholine (NEM) (543 µL, 10 eq, 4.3 mmol) was added in it. Finally compound **1** (200 mg, 1 eq, 0.43 mmol) was dissolved in dry DCM and added to the reaction mixture in drop wise manner. After that reaction mixture was left for 1.5 hr and TLC showed complete consumption of the starting material. The solvent was removed *in vacuo* and redissolved in EtOAc and washed repeatedly with water. The collected organic layer was dried over $Na_2SO_4$ and concentrated under reduced pressure. The crude product was purified by flash column chromatography using 30-40% Acetone-DCM as eluent and isolated as a pale yellow solid (405 mg, 83% yield).

**Synthesis of 1-(4,5-dimethoxy-2-nitrophenyl)ethanol (Compound 5)**

Commercially available compound 4,5-dimethoxy-2-nitrobenzaldehyde (**Compound 4**, 2 g, 9.47 mmol) was dissolved in dry DCM (20 ml) and cooled at 0°C in ice bath. Trimethylaluminimum (2 M in Toluene, 1.5 equiv, 7.1 ml, 14.2 mmol) solution was then added to it in a drop wise manner. The reaction was stirred at 0°C for 2 hrs and quenched with 1M NaOH solution and left for 30 mins more at room temperature. Then the whole reaction mixture was extracted with EtOAc and washed repeatedly with water. The collected organic layer was dried over $Na_2SO_4$ and concentrated *in vacuo*. The crude product was purified by flash column chromatography using 30% EtOAc-hexane as eluent and isolated as a pale yellow solid (1.96 g, 91% yield).

**Synthesis of ((1-(4,5-dimethoxy-2-nitrophenyl)ethoxy)methyl)(methyl)sulfane (Compound 6)**

1-(4,5-Dimethoxy-2-nitrophenyl)ethanol (**Compound 5**, 1.96 g, 8.63 mmol ) was dissolved in dry DMSO (50 equiv, 31 ml, 431.3 mmol) followed by the addition of acetic anhydride (20 equiv, 16.3 ml, 172.5 mmol) and acetic acid (20 equiv, 9.9 ml, 172.5 mmol). The reaction mixture was stirred for 24 hrs and quenched with saturated $NaHCO_3$ Solution. The reaction mixture was extracted with EtOAc and washed repeatedly with water and finally with brine. The collected organic layer was dried over $Na_2SO_4$ and concentrated *in vacuo*. The crude product was purified by column chromatography using 20% EtOAc-hexane and isolated as pale yellow solid (1.9 g, 78% yield).

**Synthesis of 1-(1-(chloromethoxy)ethyl)-4,5-dimethoxy-2-nitrobenzene (Compound 7)**

**Compound 6** (1.9 g, 6.61 mmol) was dissolved in dry DCM (15 ml) and cooled to $0^oC$ in an ice bath followed by the drop wise addition of freshly distilled $SO_2Cl_2$ (1.5 equiv, 801.5 μL, 9.92 mmol). Then the reaction was stirred for 3 hrs in ice condition. After that solvent was removed under reduced pressure in rotary evaporator and the residual $SO_2Cl_2$ was removed by repeated co-evaporation from dry benzene. The crude product was used directly for the next step without further purification.

**Synthesis of 1-((2R)-6-(((tert-butyldiphenylsilyl)oxy)methyl)-4-tritylmorpholin-2-yl)-3-((1-(4,5-dimethoxy-2-nitrophenyl)ethoxy)methyl)-5-methylpyrimidine-2,4(1H,3H)-dione (Compound 9)**

**Compound 8** (3.3 g, 4.6 mmol) was dissolved in dry DMF (10 ml) followed by the addition of $Cs_2CO_3$ (3 equiv, 4.5 g, 13.7 mmol) and cooled to $0^oC$ in ice bath. After that **compound 7** (1.5 equiv) was added in a drop wise manner dissolved in dry DMF (5 ml). The reaction mixture was left overnight and TLC showed complete consumption of the **compound 8**. Then the reaction was quenched by saturated $NH_4Cl$ and diluted with EtOAc. The organic layer was washed with water repeatedly and finally with brine. The collected organic layer was dried over $Na_2SO_4$ and concentrated *in vacuo*. The crude product was purified by column chromatography using EtOAc-hexane as eluent and isolated as a pale yellow solid (2.8 g, 65%).

**Synthesis of Compounds 10 and 11**

**Compound 10** (2.1 g, 83%, pale yellow solid) was synthesized from **compound 9** (3.4 g, 3.54 mmol) following earlier reports[1].

**Compound 11** (2 g, 81%, pale yellow solid) was synthesized from **compound 10** (2.1 g, 2.9 mmol) following earlier reports[1].

#### Spectral Data

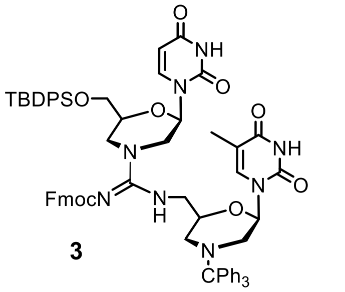


**3**

$^{1}$H NMR (300 MHz, $CDCl_3$) $\delta$ 10.40 (s, 1H), 9.98 (s, 1H), 8.66 (s, 1H), 7.79 – 7.02 (m, 35H), 6.93 (d, $J$ = 7.2 Hz, 3H), 6.31 – 6.14 (m, 1H), 5.61 (dd, $J$ = 17.3, 8.3 Hz, 2H), 4.51 (d, $J$ = 10.1 Hz, 1H), 4.28 – 4.10 (m, 2H), 3.94 (d, $J$ = 32.4 Hz, 4H), 3.76 (s, 2H), 3.52 (s, 1H), 3.19 (d, $J$ = 10.7 Hz, 2H), 2.94 (dd, $J$ = 16.6, 9.4 Hz, 2H), 2.58 (d, $J$ = 9.8 Hz, 1H),1.68 (s, 3H), 1.63 – 1.52 (m, 1H), 1.47 (d, $J$ = 10.5 Hz, 1H), 1.01 (s, 9H).

$^{13}$C NMR (75 MHz, $CDCl_3$) $\delta$ 164.4, 163.1, 150.5, 150.2, 141.3, 138.8, 135.6, 133.0, 132.8, 130.0, 129.1, 127.9, 127.8, 127.7, 127.1, 126.5, 125.5, 125.2, 120.0, 110.9, 102.9, 80.0, 77.5, 77.3, 77.1, 76.9, 76.7, 67.5, 64.1, 51.4, 50.71, 47.0, 26.8, 19.3, 12.3.

$R_f$ = 0.4 (7.5% MeOH - DCM)

HRMS (ESI) $[M + H]^{+}$: Calculated mass for $C_{70}H_{71}N_8O_9Si$ = 1195.5113 found 1195.5115.

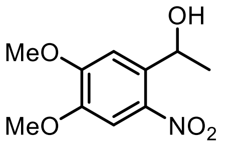


**5**

$^{1}$H NMR (300 MHz, $CDCl_3$) $\delta$ 7.52 (s, 1H), 7.30 (s, 1H), 5.53 (q, $J$ = 6.3 Hz, 1H), 3.99 (s, 3H), 3.93 (s, 3H), 2.88 (s, 1H), 1.53 (d, $J$ = 6.4 Hz, 3H).

$^{13}$C NMR (75 MHz, $CDCl_3$) $\delta$ 153.7, 147.6, 139.5, 137.1, 108.5, 107.6, 65.7, 56.4, 56.3, 24.4.

HRMS (ESI) $[M + Na]^{+}$: Calculated mass for $C_{10}H_{13}NO_5Na$ = 250.0691, found 250.0692.

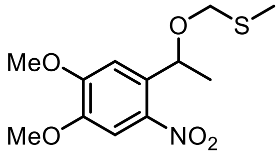


**6**

$^{1}$H NMR (300 MHz, $CDCl_3$) $\delta$ 7.47 (s, 1H), 7.12 (s, 1H), 5.42 (q, $J$ = 6.2 Hz, 1H), 4.52 (d, $J$ = 11.4 Hz, 1H), 4.24 (d, $J$ = 11.4 Hz, 1H), 3.90 (s, 3H), 3.84 (s, 3H), 2.03 (s, 3H), 1.42 (d, $J$ = 6.3 Hz, 3H).

$^{13}$C NMR (75 MHz, $CDCl_3$) $\delta$ 153.6, 147.6, 140.2, 134.3, 108.4, 107.4, 77.4, 72.9, 70.3, 56.2, 56.0, 23.1, 13.9.

HRMS (ESI) $[M + Na]^{+}$: Calculated mass for $C_{12}H_{17}NO_5SNa$ = 310.0725, found 310.0724.

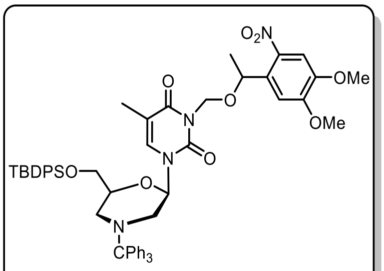


9

$^1$H NMR (300 MHz, CDCl$_3$) $\delta$ 7.65 – 7.44 (m, 11H), 7.44 – 7.28 (m, 13H), 7.22 (d, $J$ = 6.4 Hz, 4H), 6.78 (dd, $J$ = 9.9, 1.3 Hz, 1H), 6.04 (ddd, $J$ = 16.3, 9.7, 2.3 Hz, 1H), 5.57 – 5.35 (m, 2H), 5.26 (dd, $J$ = 16.8, 9.4 Hz, 1H), 4.29 (d, $J$ = 12.9 Hz, 1H), 4.01 (s, 1H), 3.92 – 3.69 (m, 6H), 3.58 (ddd, $J$ = 10.5, 6.2, 3.0 Hz, 1H), 3.39 – 3.20 (m, 2H), 1.68 (dd, $J$ = 7.8, 1.2 Hz, 3H), 1.55 (dd, $J$ = 6.2, 1.0 Hz, 4H), 0.97 (d, $J$ = 3.6 Hz, 9H).

$^{13}$C NMR (75 MHz, CDCl$_3$) δ 162.9, 153.6, 153.5, 150.5, 150.3, 147.7, 147.6, 140.2, 140.0, 135.8, 135.6, 135.5, 135.4, 134.5, 134.3, 133.4, 133.3, 133.2, 133.1, 129.9, 129.8, 129.7, 129.6, 129.3, 127.9, 127.8, 127.7, 127.6, 126.6, 126.5, 109.6, 109.4, 108.9, 108.7, 107.4, 81.1, 80.9, 77.4, 77.1, 77.0, 76.9, 76.8, 73.5, 73.4, 70.2, 64.6, 64.5, 56.6, 56.4, 56.2, 52.1, 51.7, 50.1, 49.9, 26.8, 23.9, 19.3, 12.9, 12.8.

HRMS (ESI) [M + Na]$^+$:Calculated mass for $C_{56}H_{60}N_4O_9SiNa$ = 983.4027, found 983.4026.

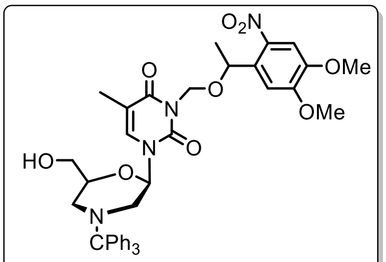


10 (US)

$^1$H NMR (300 MHz, CDCl$_3$) $\delta$ 7.46 (d, $J$ = 7.7 Hz, 6H), 7.37 (s, 1H), 7.35 – 7.24 (m, 8H), 7.19 (dd, $J$ = 14.7, 7.4 Hz, 4H), 6.62 (d, $J$ = 1.4 Hz, 1H), 6.09 (dd, $J$ = 9.7, 2.4 Hz, 1H), 5.57 (dd, $J$ = 10.0, 2.4 Hz, 1H), 5.44 – 5.30 (m, 1H), 5.22 (d, $J$ = 9.9 Hz, 1H), 4.24 (ddd, $J$ = 9.0, 5.3, 2.5 Hz, 1H), 4.02 (s, 3H), 3.75 (d, $J$ = 1.6 Hz, 3H), 3.54 (q, $J$ = 5.2, 4.2 Hz, 2H), 3.17 (dt, $J$ = 11.3, 2.4 Hz, 2H), 3.06 (dt, $J$ = 11.9, 2.3 Hz, 1H), 1.65 (d, $J$ = 1.6 Hz, 3H), 1.55 (d, $J$ = 6.3 Hz, 3H), 1.47 – 1.27 (m, 2H).

$^{13}$C NMR (75 MHz, CDCl$_3$) δ 162.8, 153.1, 150.1, 147.3, 139.6, 136.4, 133.9, 129.2, 127.9, 127.8, 126.6, 109.9, 108.7, 106.7, 80.2, 78.5, 77.4, 76.8, 73.9, 70.5, 64.1, 56.3, 56.2, 51.6, 48.6, 23.9, 12.7.

$R_f$ = 0.5 (5% Acetone- DCM)

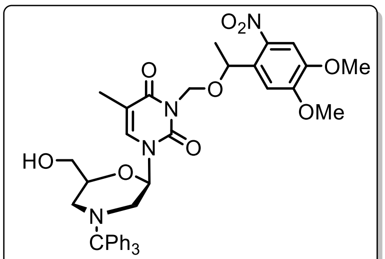


10 (LS)

$^1$H NMR (300 MHz, CDCl$_3$) δ 7.67 – 7.37 (m, 7H), 7.37 – 7.24 (m, 8H), 7.24 – 7.10 (m, 4H), 6.81 (t, $J$ = 1.4 Hz, 1H), 6.03 (dd, $J$ = 9.7, 2.3 Hz, 1H), 5.51 – 5.34 (m, 2H), 5.27 (d, $J$ = 9.7 Hz, 1H), 4.26 (ddd, $J$ = 6.7, 4.6, 2.5 Hz,

1H), 3.89 (d, *J* = 1.6 Hz, 3H), 3.86 – 3.79 (m, 3H), 3.57 (q, *J* = 6.2 Hz, 2H), 3.26 (dt, *J* = 11.3, 2.4 Hz, 1H), 3.10 (d, *J* = 11.8 Hz, 1H), 2.15 (s, 1H), 1.72 (t, *J* = 1.5 Hz, 3H), 1.53 (d, *J* = 6.2 Hz, 3H), 1.50 – 1.29 (m, 2H).

$^{13}$C NMR (75 MHz, $CDCl_3$) δ 162.9, 153.6, 150.3, 147.5, 140.1, 135.9, 134.3, 129.2, 127.9, 126.5, 109.7, 108.9, 107.8, 81.4, 77.7, 77.4, 76.8, 73.6, 70.2, 63.7, 56.6, 56.4, 51.8, 48.9, 23.9, 12.9.

$R_f$ = 0.4 (5% Acetone- DCM)

HRMS (ESI) $[M + Na]^+$: Calculated mass for $C_{40}H_{42}N_4O_9Na$ = 745.2849, found 745.2851.

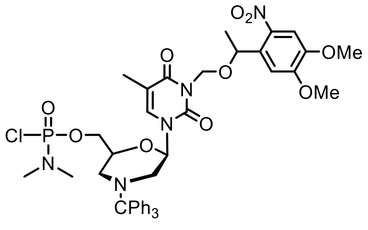


11(US)

$^1$H NMR (300 MHz, $CDCl_3$) *δ* 7.44 (d, *J* = 6.1 Hz, 7H), 7.38 – 7.24 (m, 8H), 7.18 (t, *J* = 7.2 Hz, 3H), 6.85 (dd, *J* = 3.3, 1.3 Hz, 1H), 6.07 (dt, *J* = 9.5, 2.0 Hz, 1H), 5.48 – 5.32 (m, 2H), 5.18 (d, *J* = 9.4 Hz, 1H), 4.45 – 4.29 (m, 1H), 4.08 (dddd, *J* = 12.9, 9.2, 5.2, 2.9 Hz, 2H), 3.98 (s, 3H), 3.75 (s, 3H), 3.26 (dt, *J* = 11.2, 2.4 Hz, 1H), 3.16 (dt, *J* = 11.9, 2.3 Hz, 1H), 2.61 (dd, *J* = 13.9, 2.4 Hz, 6H), 1.79 – 1.69 (m, 3H), 1.52 (s, 2H), 1.50 – 1.33 (m, 3H).

$^{13}$C NMR (75 MHz, $CDCl_3$) *δ* 162.8, 153.6, 150.4, 147.6, 140.1, 135.4, 134.2, 134.1, 129.2, 128.0, 127.9, 127.8, 126.6, 109.9, 108.7, 107.2, 80.8, 77.4, 76.9, 74.6, 74.5, 74.4, 73.3, 70.1, 67.2, 67.1, 56.5, 56.2, 51.4, 49.0, 36.7, 36.6, 36.5, 23.8, 12.9.

$^{31}$P NMR (121 MHz, $CDCl_3$) δ 18.29, 18.00.

$R_f$ = 0.7 (5% Acetone- DCM)

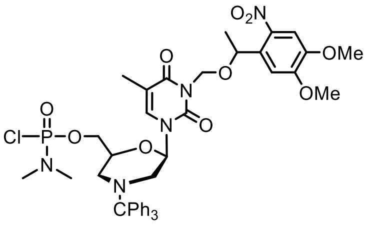


11(LS)

$^1$H NMR (300 MHz, $CDCl_3$) *δ* 7.50 (dd, *J* = 19.7, 4.5 Hz, 7H), 7.38 – 7.25 (m, 9H), 7.25 – 7.12 (m, 5H), 6.91 – 6.76 (m, 1H), 6.04 (dd, *J* = 9.6, 2.4 Hz, 1H), 5.48 – 5.31 (m, 2H), 5.24 (d, *J* = 9.3 Hz, 1H), 4.40 (ddt, *J* = 7.5, 4.5, 2.5 Hz, 1H), 4.18 – 4.02 (m, 2H), 3.91 (d, *J* = 1.3 Hz, 3H), 3.84 (s, 3H), 3.29 (dt, *J* = 11.4, 2.1 Hz, 1H), 3.24 – 3.11 (m, 1H), 2.64 (dd, *J* = 13.9, 1.0 Hz, 6H), 1.73 (dd, *J* = 2.9, 1.2 Hz, 4H), 1.53 (dd, *J* = 6.3, 0.9 Hz, 4H).

$^{13}C$ NMR (75 MHz, $CDCl_3$) $\delta$ 162.9, 162.8, 153.6, 150.4, 147.7, 140.1, 135.7, 135.6, 134.2, 134.1, 129.3, 128.1, 127.9, 126.7, 109.9, 109.0, 108.9, 107.4, 81.1, 77.4, 76.9, 74.9, 74.8, 73.3, 70.1, 67.3, 67.2, 56.6, 56.5, 56.4, 51.7, 49.1, 36.8, 36.6, 23.8, 12.9, 12.8.

$^{31}P$ NMR (121 MHz, $CDCl_3$) δ 18.35, 18.08.

$R_f$ = 0.6 (5% Acetone- DCM)

HRMS (ESI) $[M + Na]^+$: Calculated mass for $C_{42}H_{47}N_4O_{10}PClNa$ = 870.2647 found 870.2645.

#### NMR Spectra

[1]H NMR (300 MHz, $CDCl_3$) of **Compound 3**

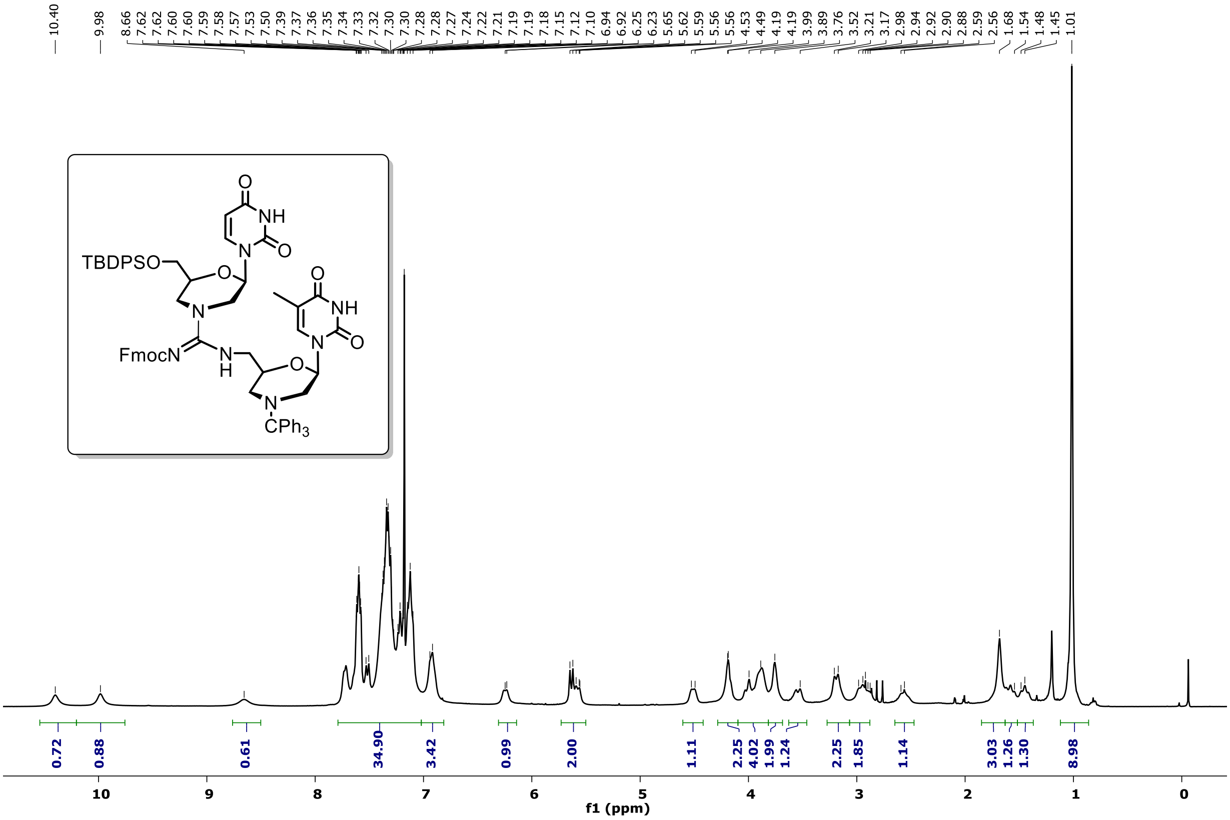


[13]C NMR (75 MHz, $CDCl_3$) of **Compound 3**

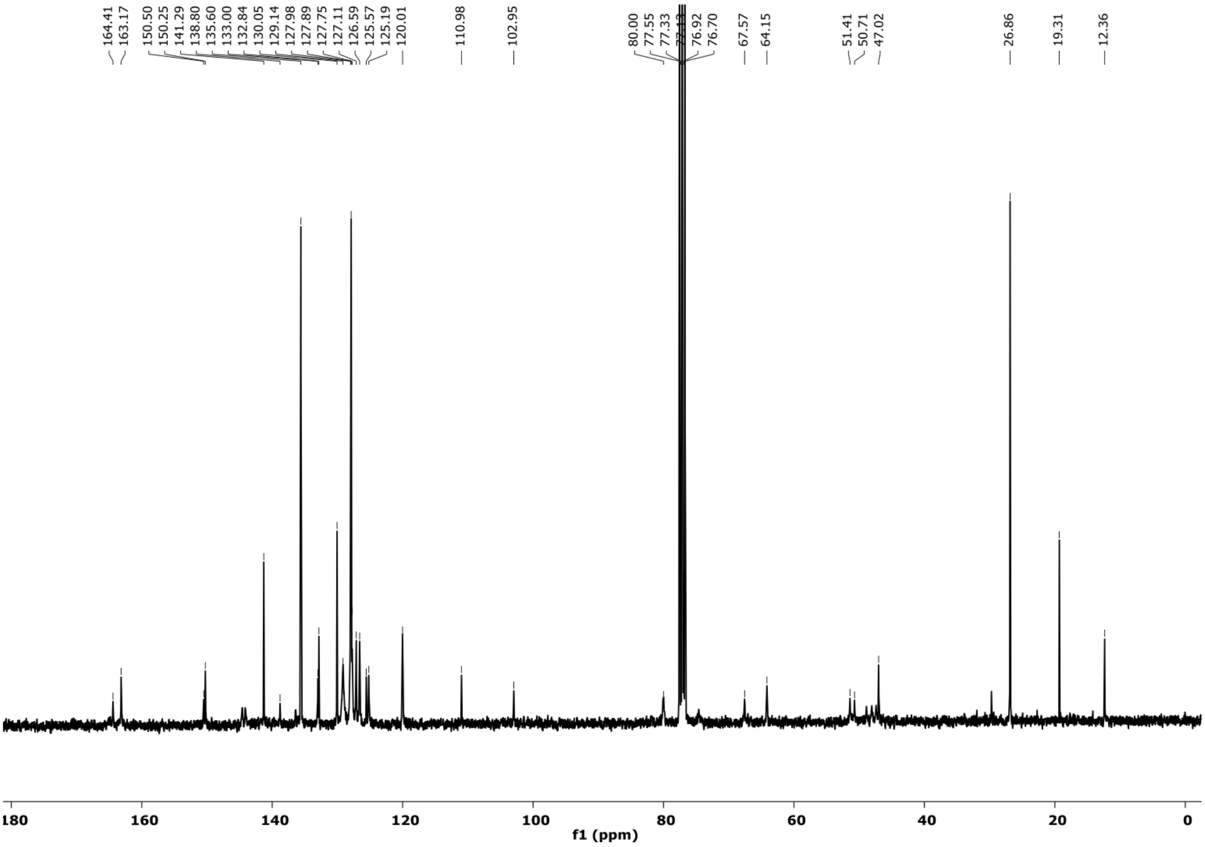

$^1$H NMR (300 MHz, $CDCl_3$) of **Compound 5**

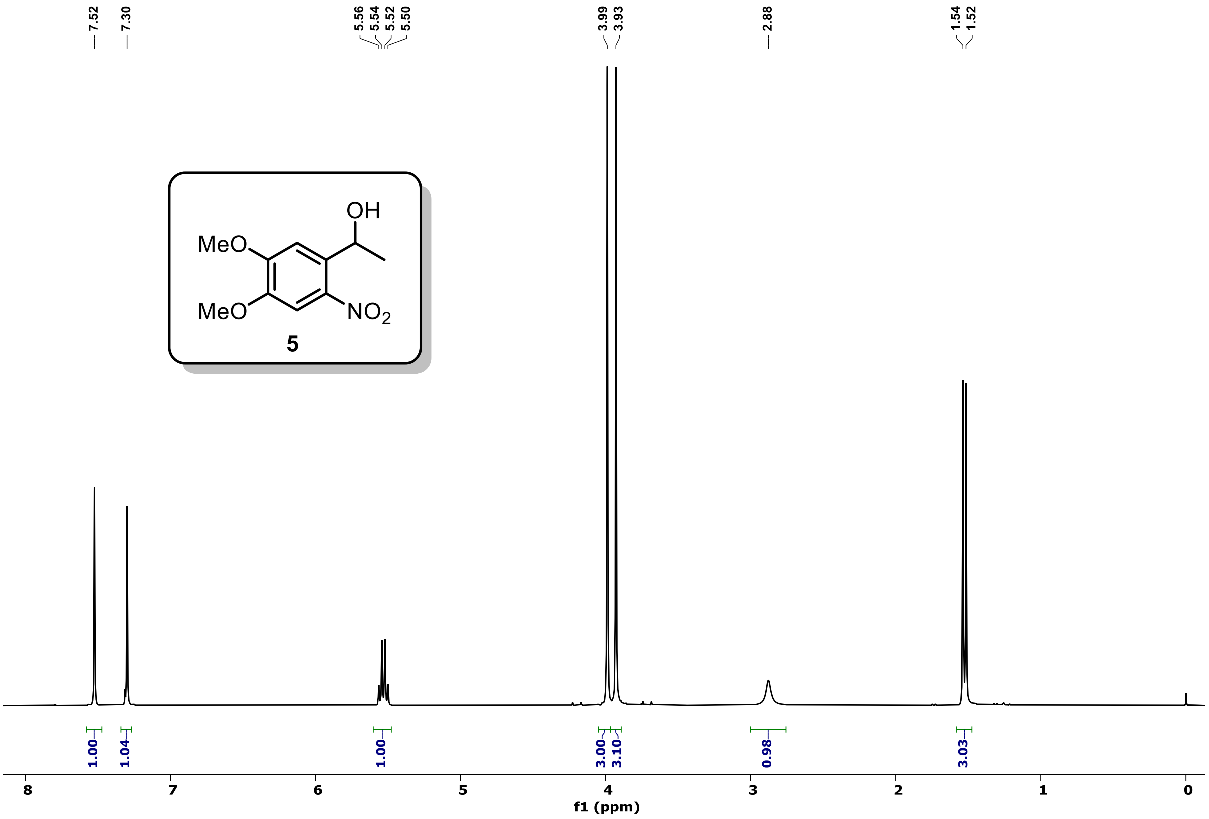


$^{13}$C NMR (75 MHz, $CDCl_3$) of **Compound 5**

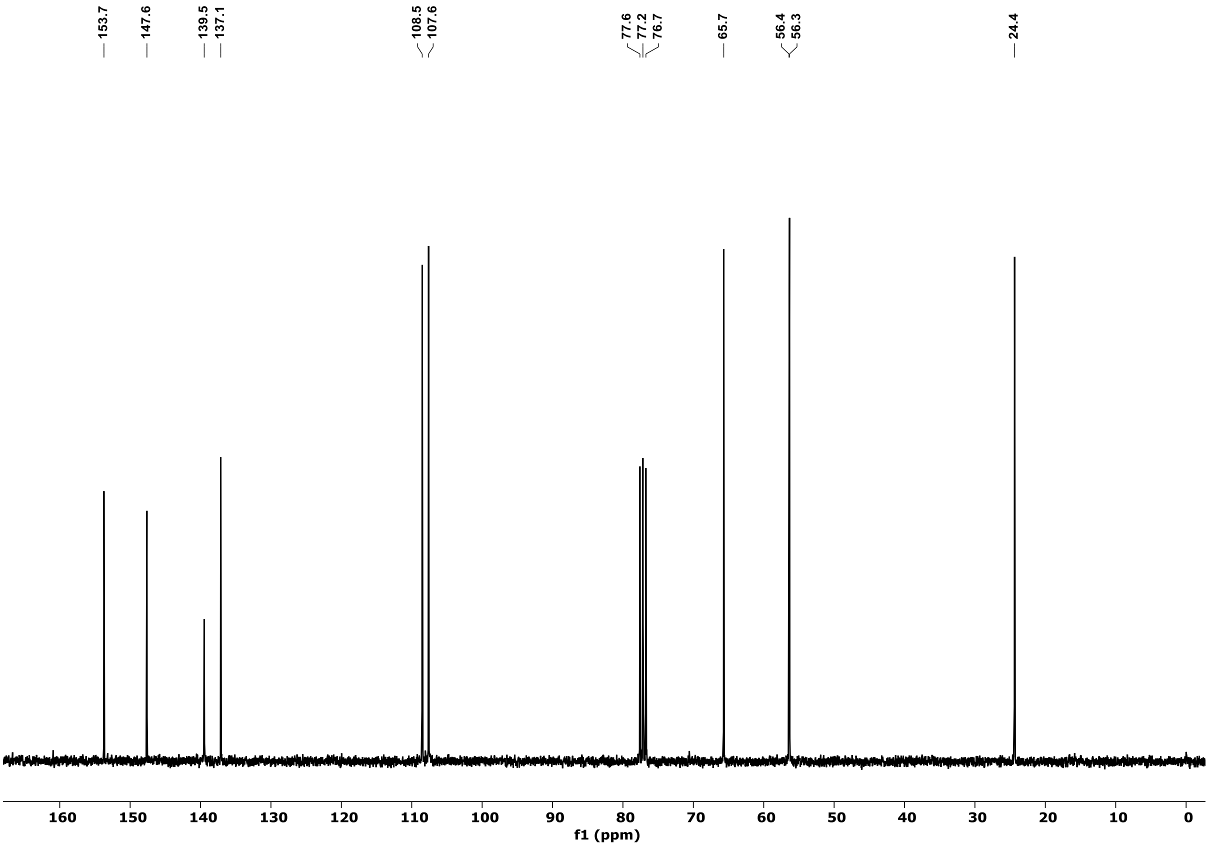

### $^{1}$H NMR (300 MHz, CDCl$_3$) of **Compound 6**

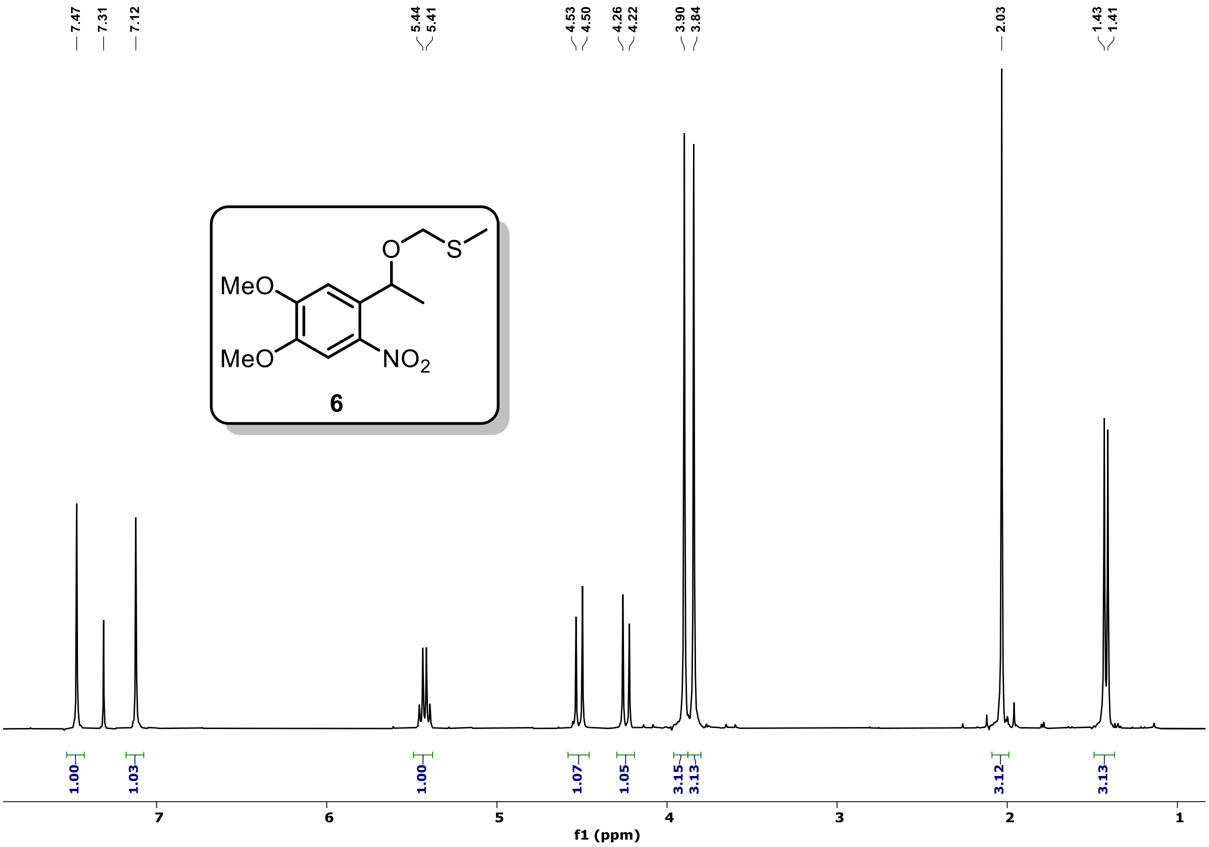


### $^{13}$C NMR (75 MHz, CDCl$_3$) of **Compound 6**

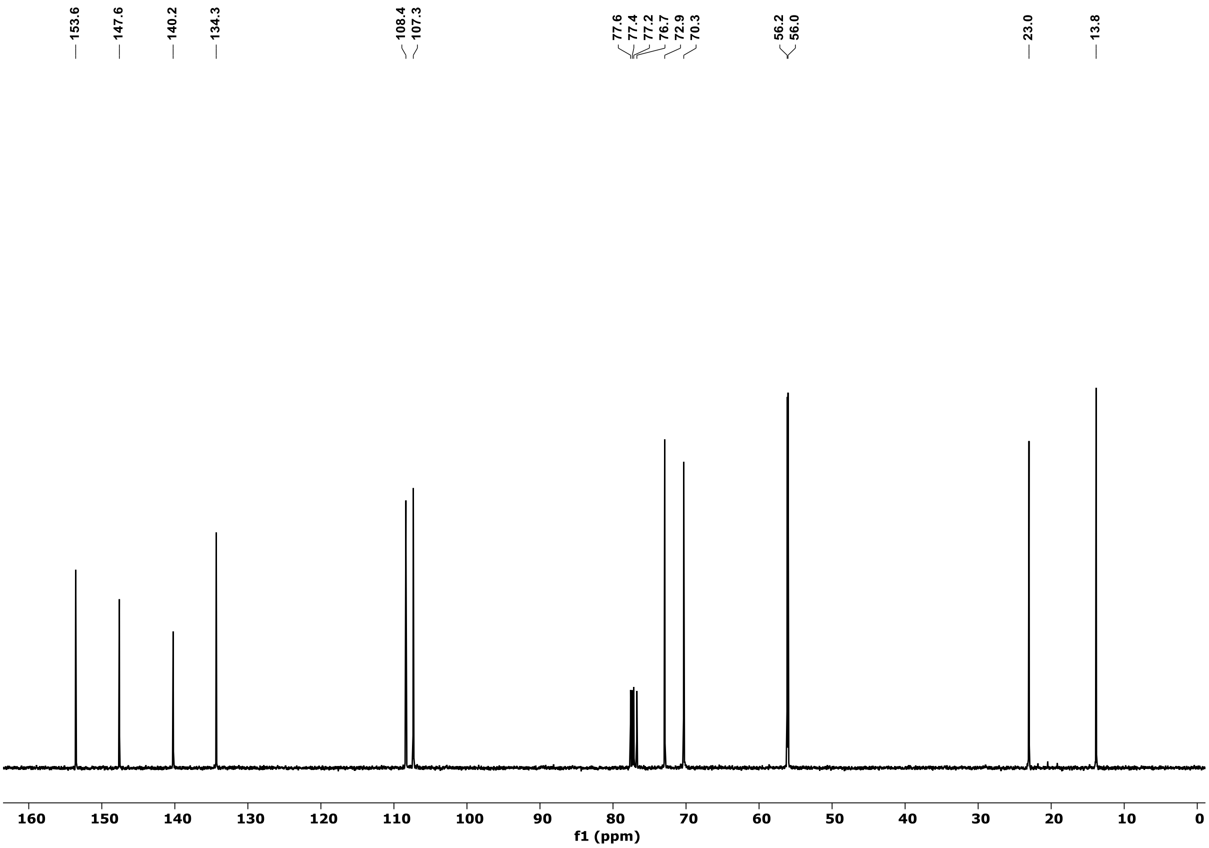

### $^{1}H$ NMR (300 MHz, $CDCl_3$) of **Compound 9**

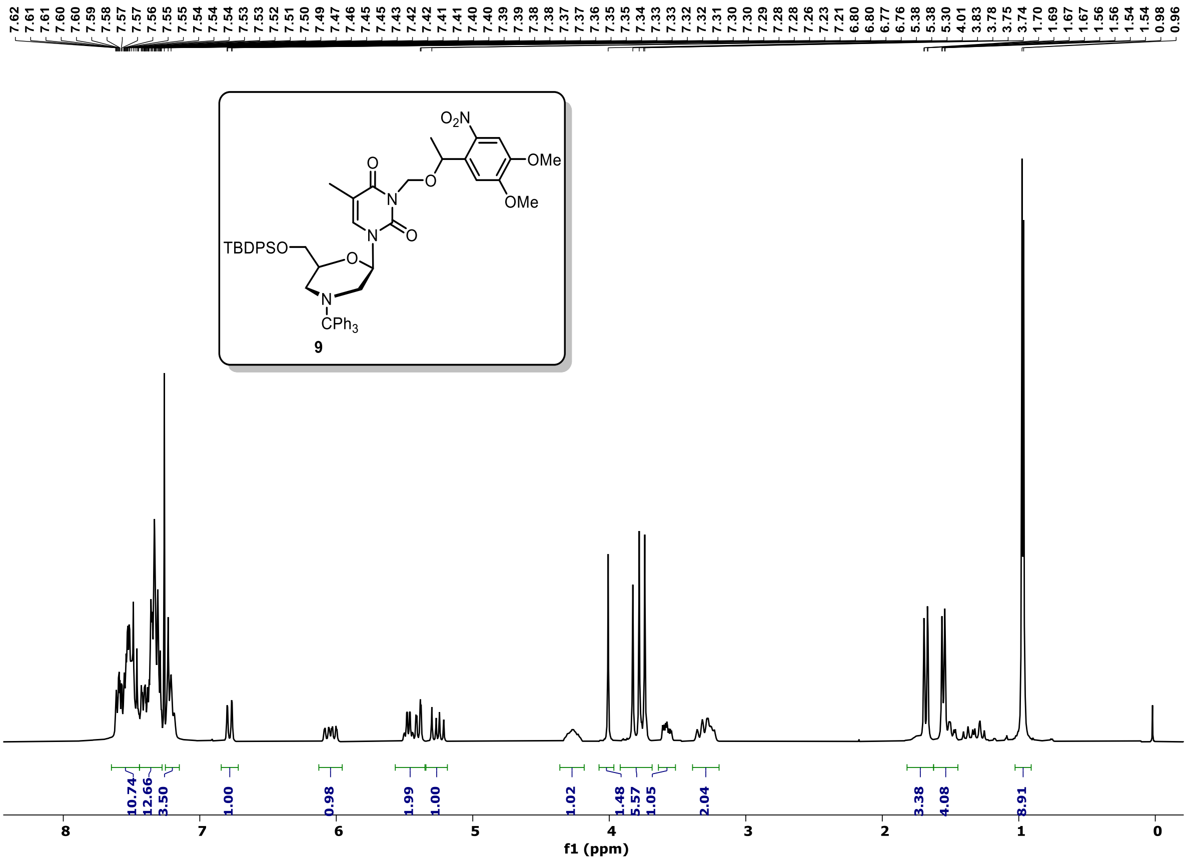


### $^{13}C$ NMR (75 MHz, $CDCl_3$) of **Compound 9**

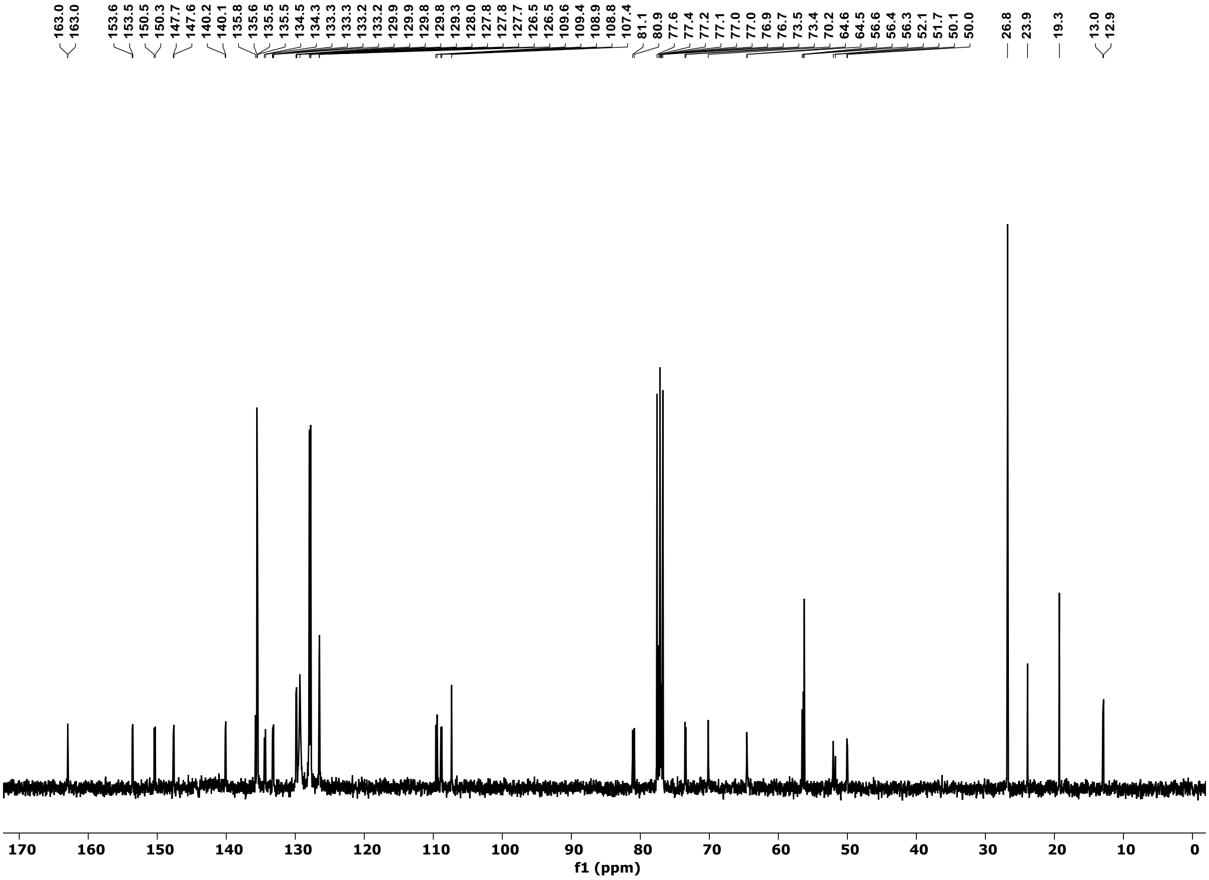

<sup></sup>

$^{1}$H NMR (300 MHz, $CDCl_3$) of **Compound 10 (US)**

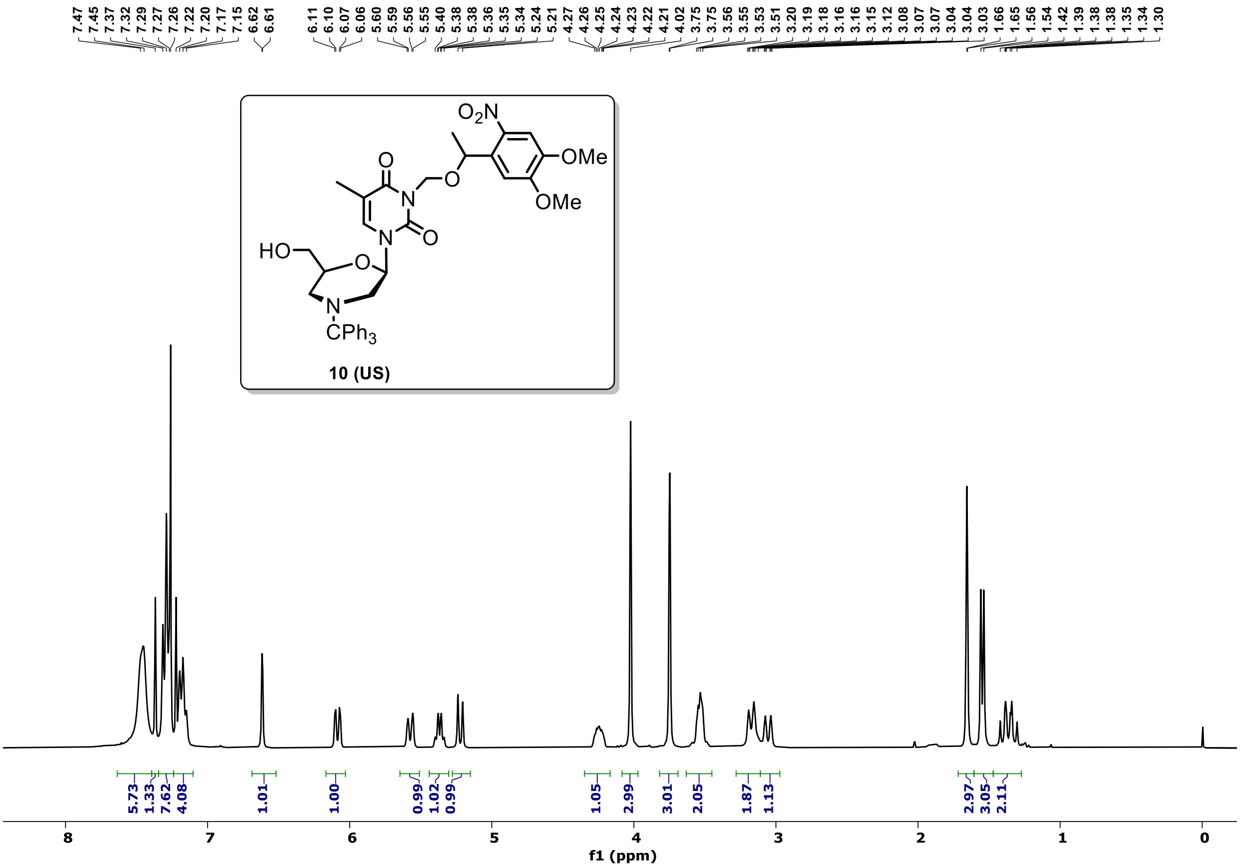


$^{13}$C NMR (75 MHz, $CDCl_3$) of **Compound 10 (US)**

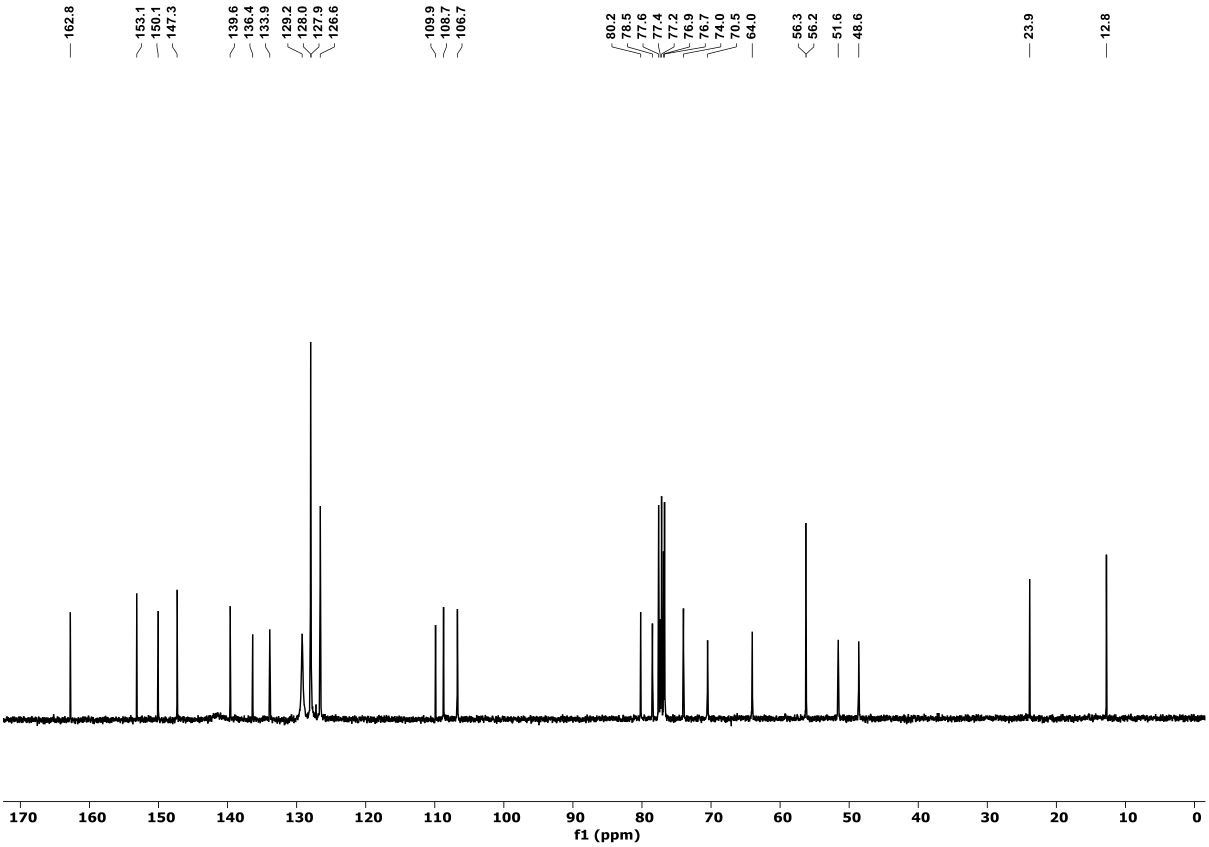

$^1$H NMR (300 MHz, CDCl$_3$) of **Compound 10 (LS)**

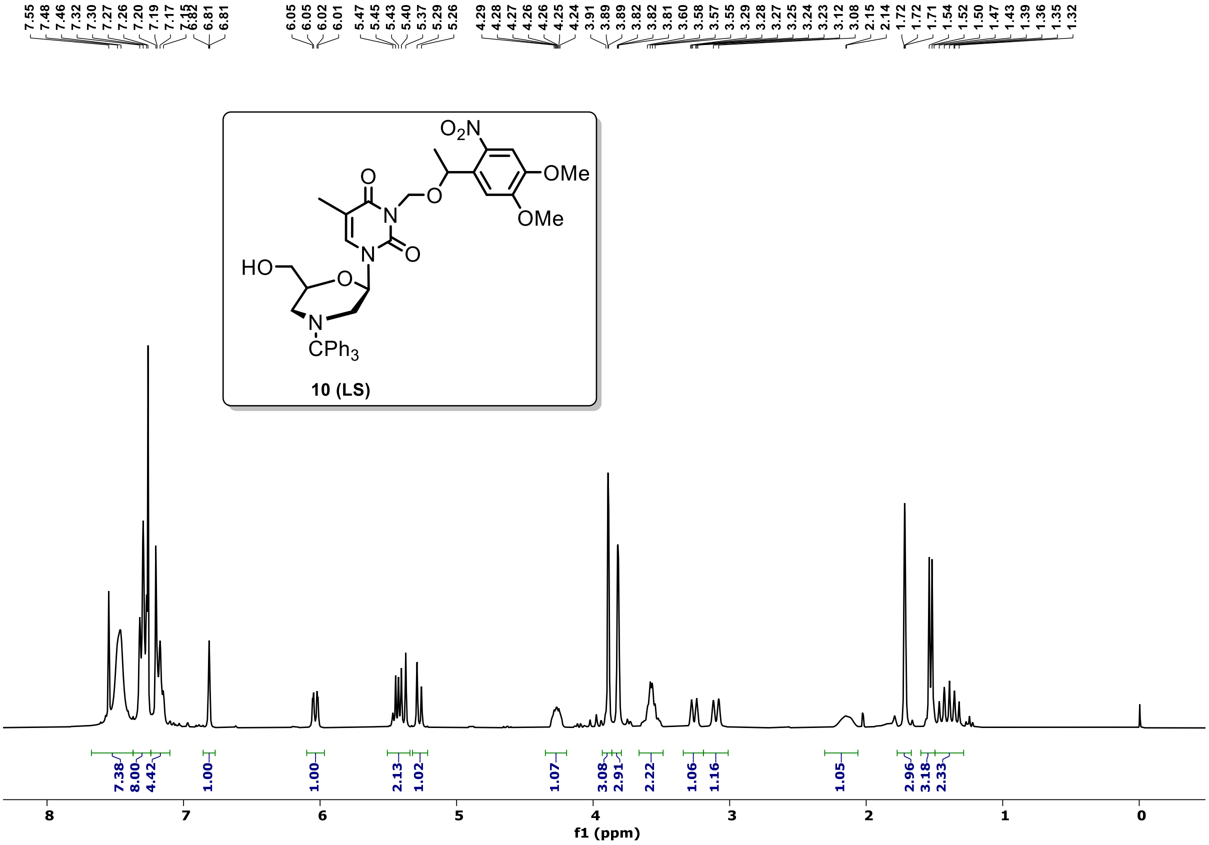


$^{13}$C NMR (75 MHz, CDCl$_3$) of **Compound 10 (LS)**

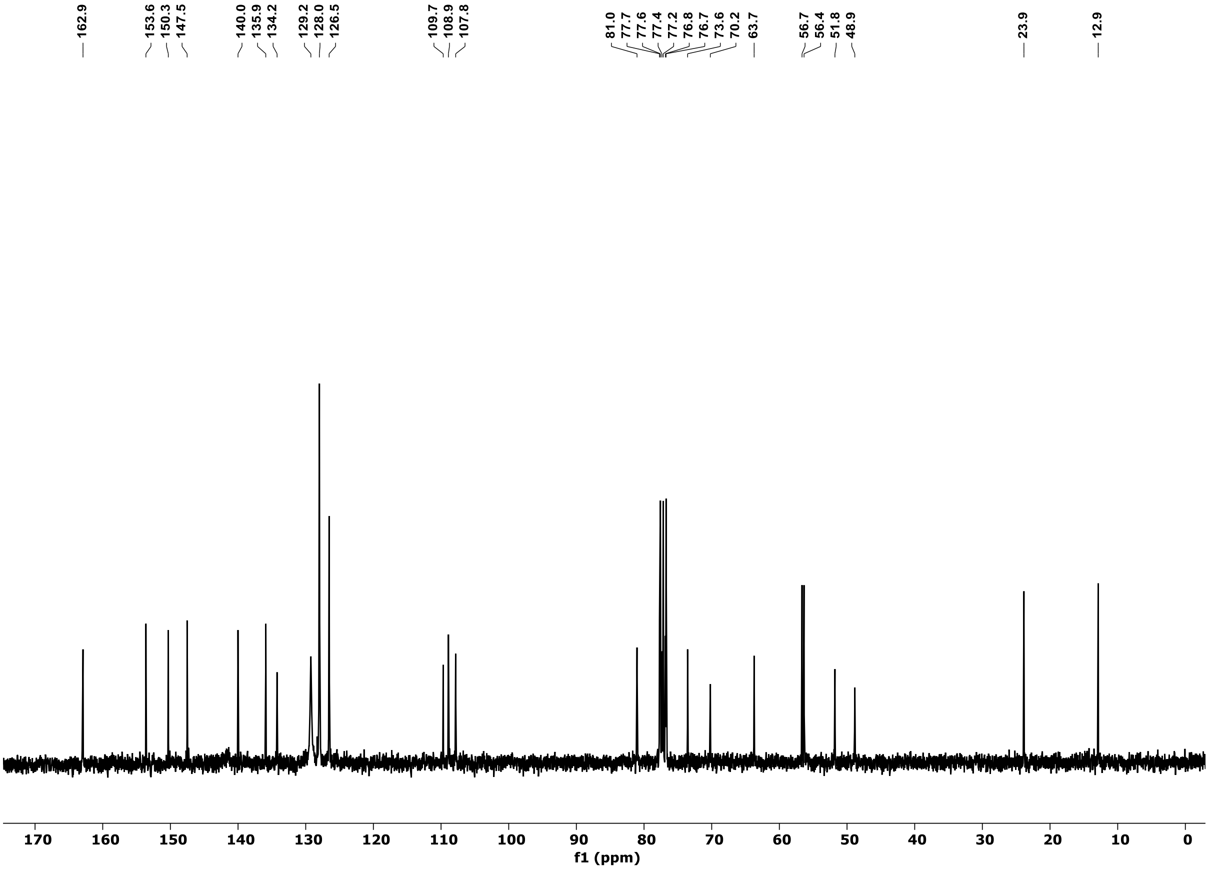

$^{1}H$ NMR (300 MHz, $CDCl_3$) of **Compound 11 (US)**

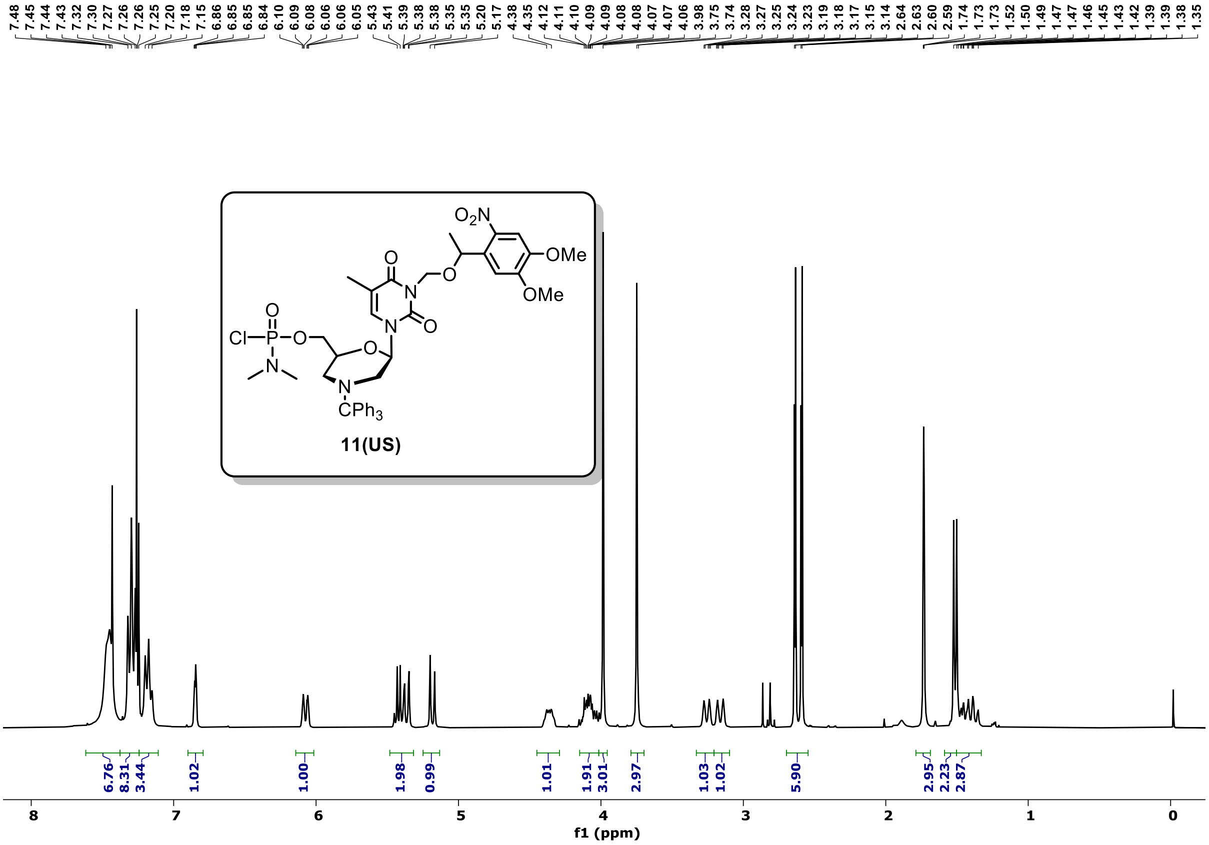


$^{13}C$ NMR (75 MHz, $CDCl_3$) of **Compound 11 (US)**

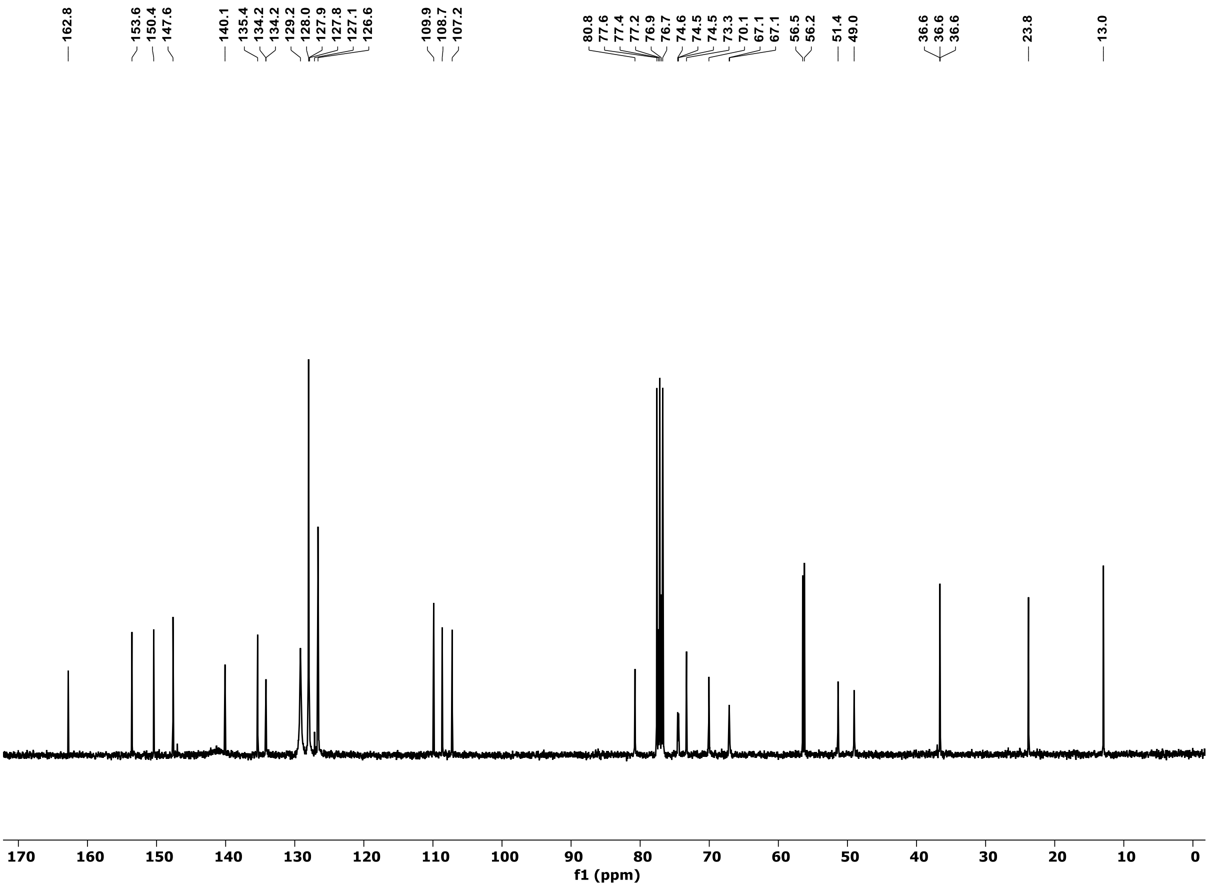

$^{31}P$ NMR (121 MHz, $CDCl_3$) of **Compound 11 (US)**

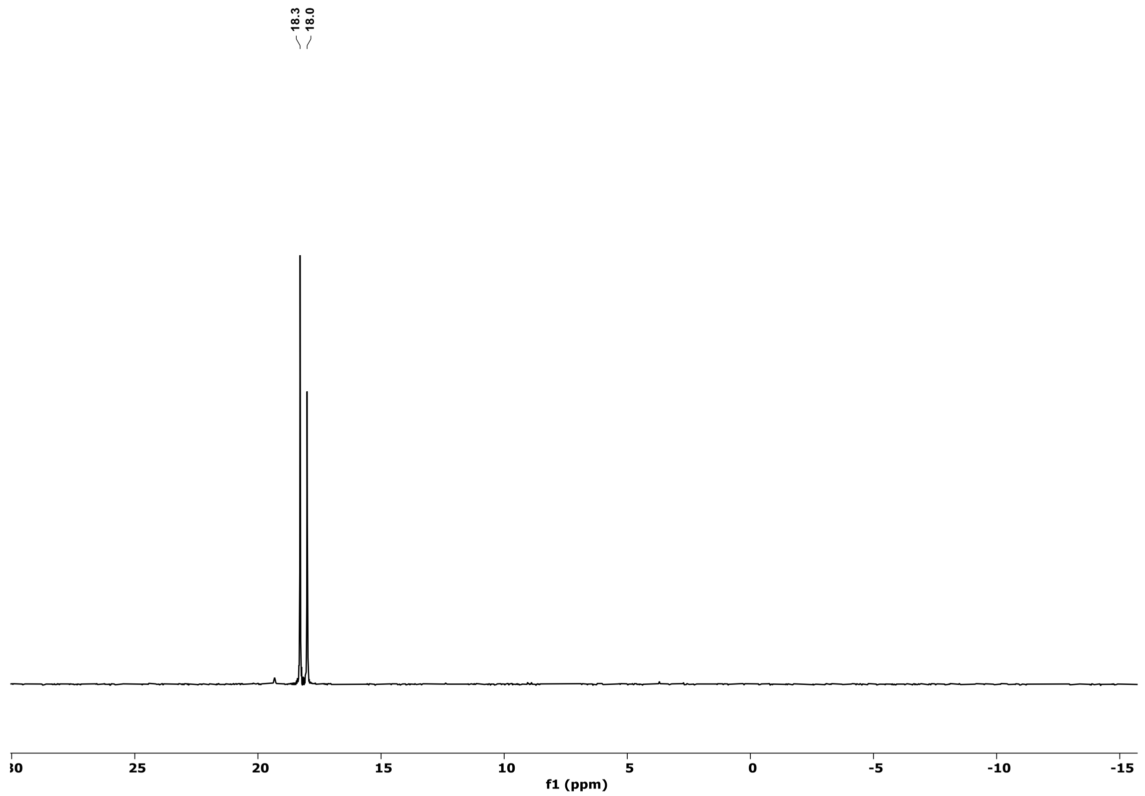


$^{1}H$ NMR (300 MHz, $CDCl_3$) of **Compound 11 (LS)**

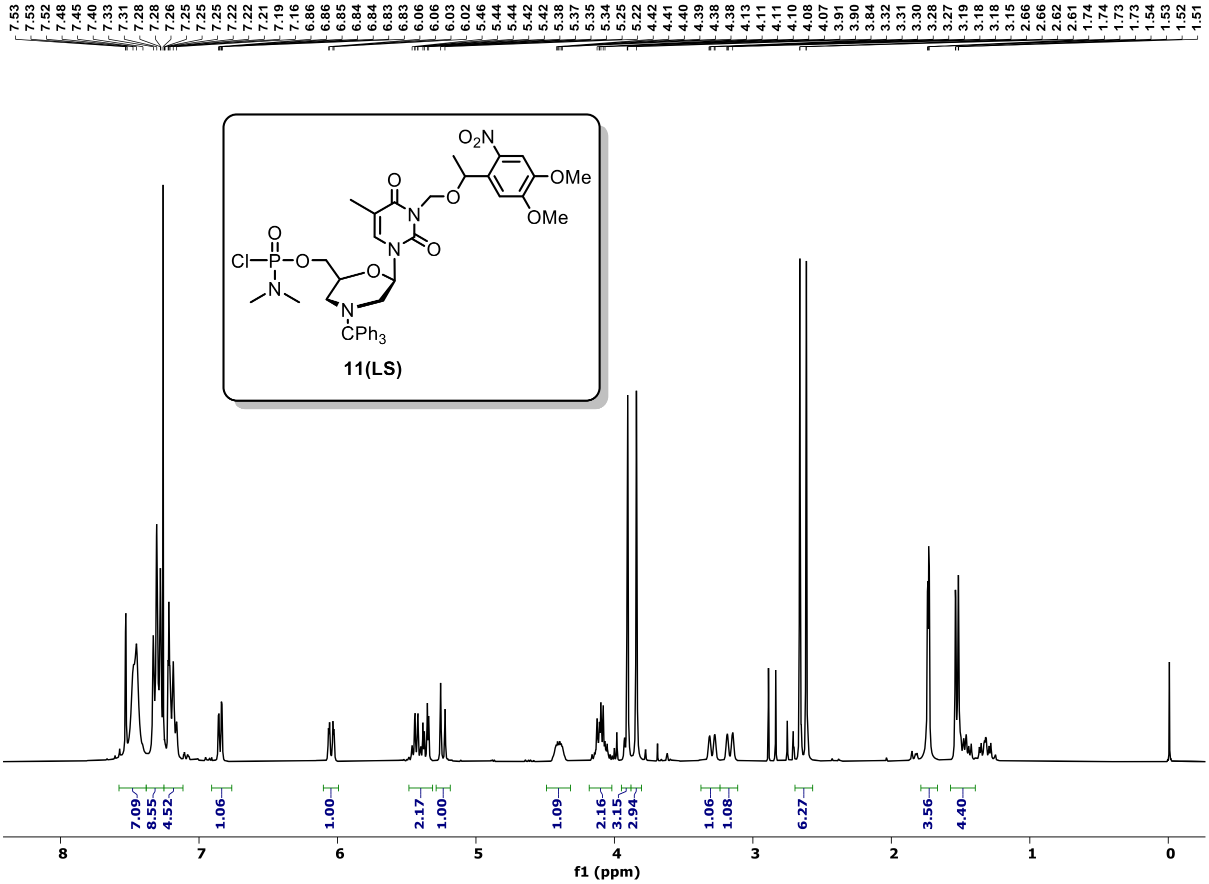

#### $^{13}C$ NMR (75 MHz, $CDCl_3$) of **Compound 11 (LS)**

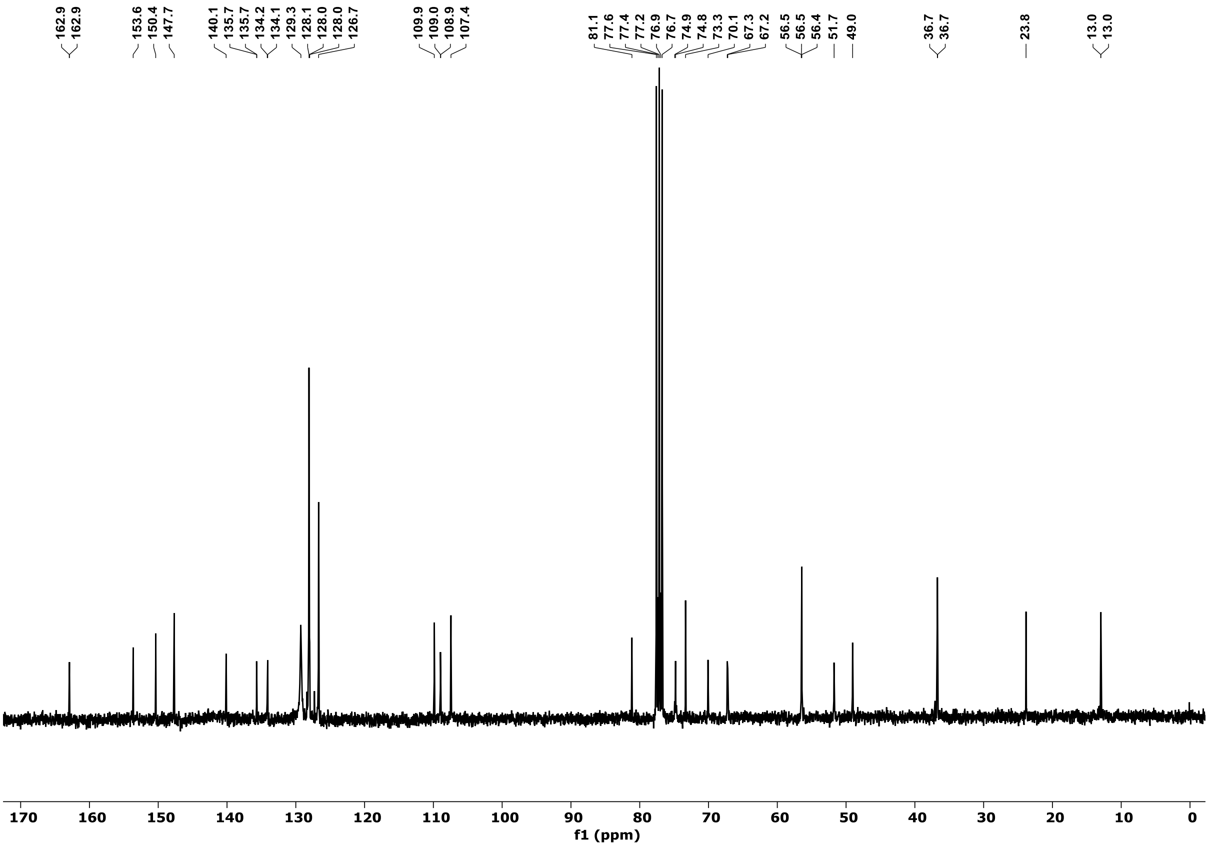


#### $^{31}P$ NMR (121 MHz, $CDCl_3$) of **Compound 11 (LS)**

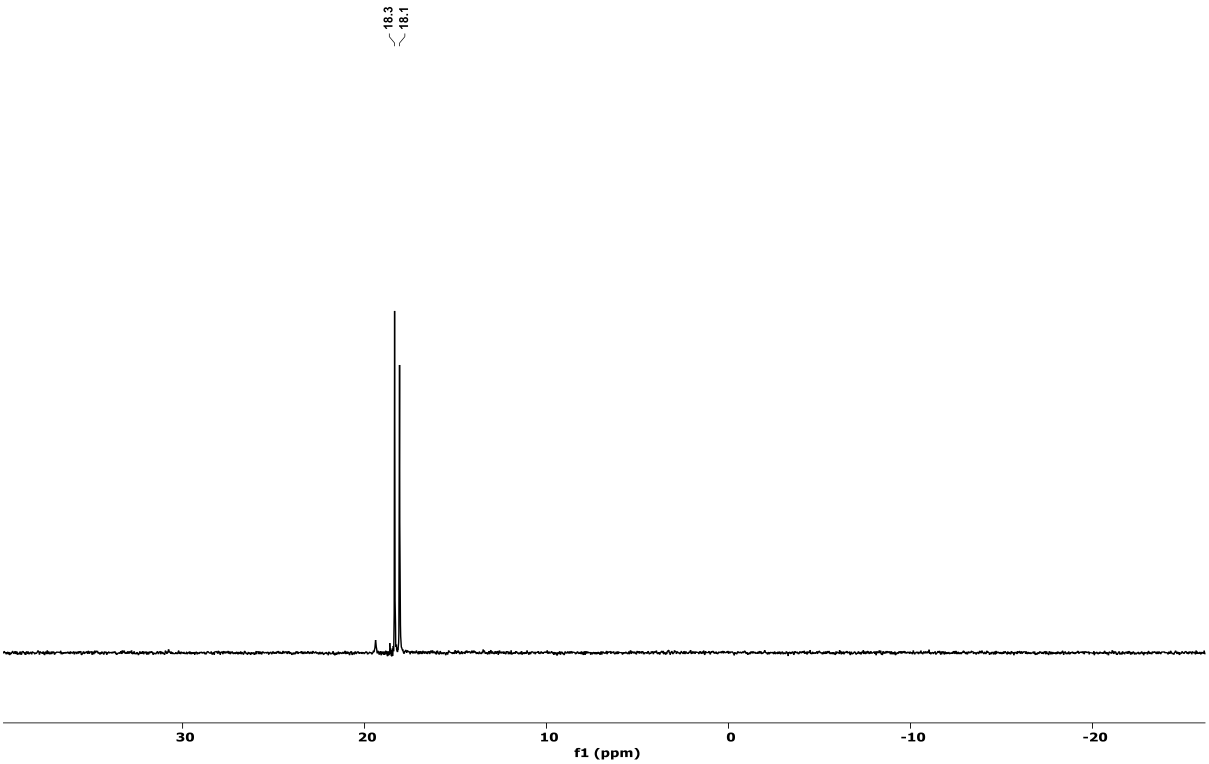

**Table S3**. HPLC retention time of PMOs

| Sequence | Retention time($R_t$) | |
|---|---|---|
| 5′-T^T^T^T^TTTTTT-Tr-3′ (GMOPMO-1) | 23.5 | |
| 5′-T^A^T^A^ATTTTT-Tr-3′ (GMOPMO-2) | 23.3 | |
| **Sequence** | **Before irradiation** | **After irradiation** |
| 5'-TATAAATTGTAACTGAGGTAAGAGG-3' (cPMO1) | 19.1 | 15.1 |
| 5'-T^A^T^A^AATTGTAACTGAGGTAAGAGG-3'(cPMO2) | 19.1 | 14.5 |
| **Sequence** | **Retention time($R_t$)** | |
| 5′-T^A^T^A^AATTGTAACTGAGGTAAGAGG-3′(PMO 3) | 15.3 | |
| 5'-CCTCTTACCTCAGTTACAATTTATA-3' (tbMO) | 13.2 | |
| PMO1-BODIPY | 21.9 | |
| PMO2-BODIPY | 20.1 | |
| PMO3-BODIPY | 20.2 | |

**HPLC Chromatogram of PMOs:**

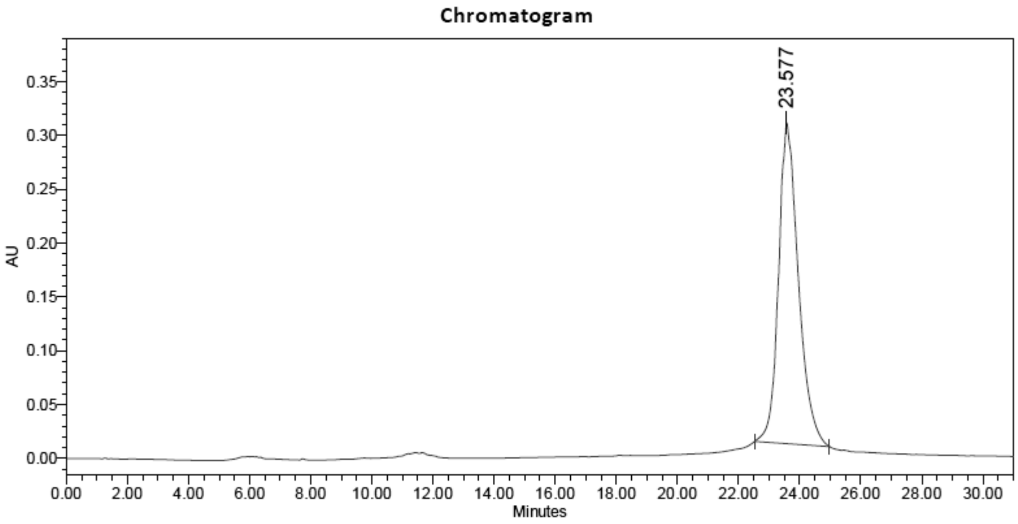


Peak table

| | RT | Area | Height | % Area |
|---|---|---|---|---|
| 1 | 23.577 | 13438568 | 297966 | 100.00 |

**Figure S13.** HPLC chromatogram of complimentary GMOPMO-1 (5′-T^T^T^T^TTTTTT-Tr-3′) (Trityl ON mode)

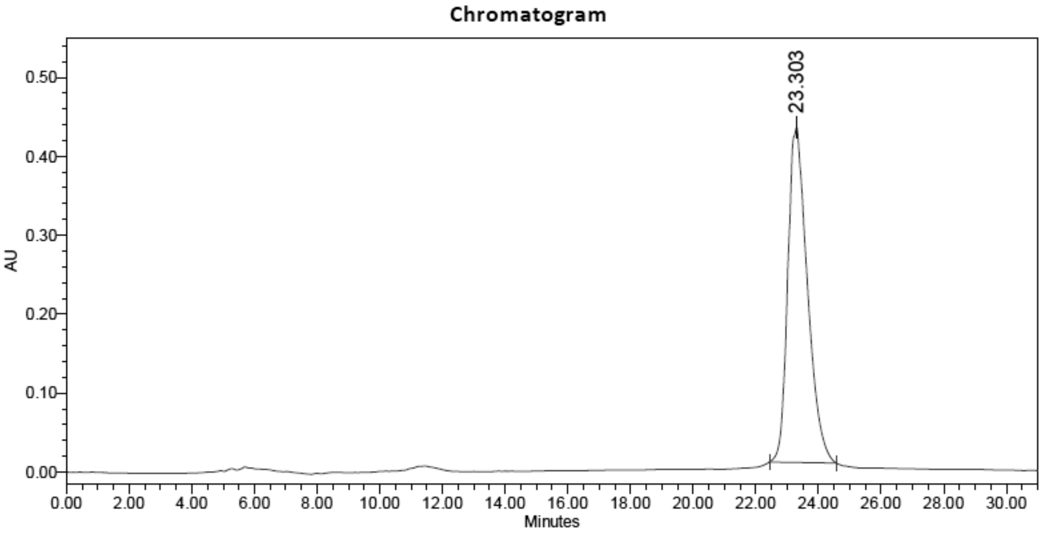


Peak table

| | RT | Area | Height | % Area |
|---|---|---|---|---|
| 1 | 23.303 | 18459254 | 424913 | 100.00 |

**Figure S14.** HPLC chromatogram of complimentary GMOPMO-2 (5′-T^A^T^A^ATTTTT-Tr-3′) (Trityl On mode)

**Chromatogram**

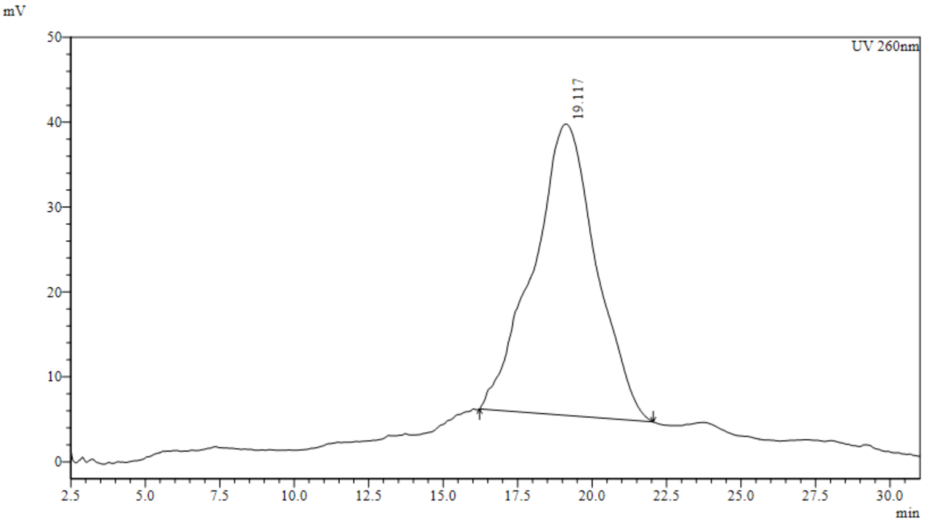


**Peak Table**

UV 260nm

| Peak# | Ret. Time | Area | Height | Area% |
|---|---|---|---|---|
| 1 | 19.117 | 5059449 | 34305 | 100.000 |
| Total | | 5059449 | 34305 | 100.000 |

**Figure S15.** HPLC chromatogram of cPMO1 (5′-TATAAATTGTAACTGAGGTAAGAGG-3′)

**Chromatogram**

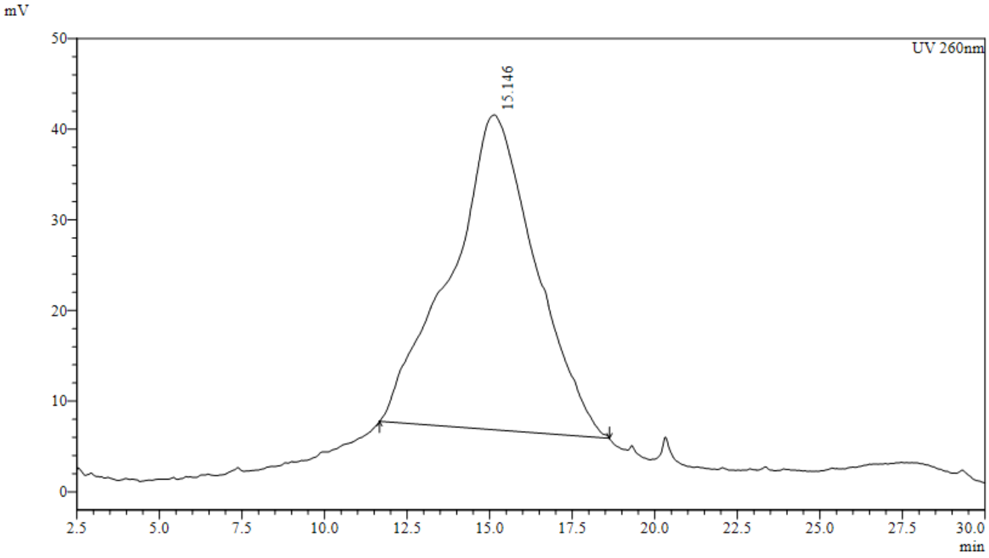


**Peak Table**

UV 260nm

| Peak# | Ret. Time | Area | Height | Area% |
|---|---|---|---|---|
| 1 | 15.146 | 6227758 | 34755 | 100.000 |
| Total | | 6227758 | 34755 | 100.000 |

**Figure S16.** HPLC chromatogram of PMO1 (after irradiation) (5′-TATAAATTGTAACTGAGGTAAGAGG-3′)

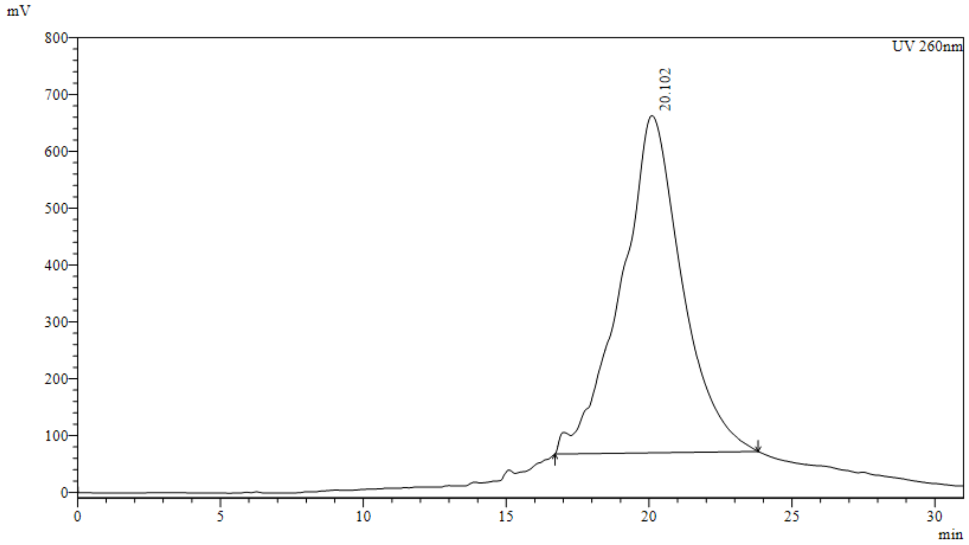


**Peak Table**

UV 260nm

| Peak# | Ret. Time | Area | Height | Area% |
|---|---|---|---|---|
| 1 | 20.102 | 88143669 | 592923 | 100.000 |
| Total | | 88143669 | 592923 | 100.000 |

**Figure S17.** HPLC chromatogram of cPMO2 (5′-T^A^T^A^AATTGTAACTGAGGTAAGAGG-3′)

#### Chromatogram

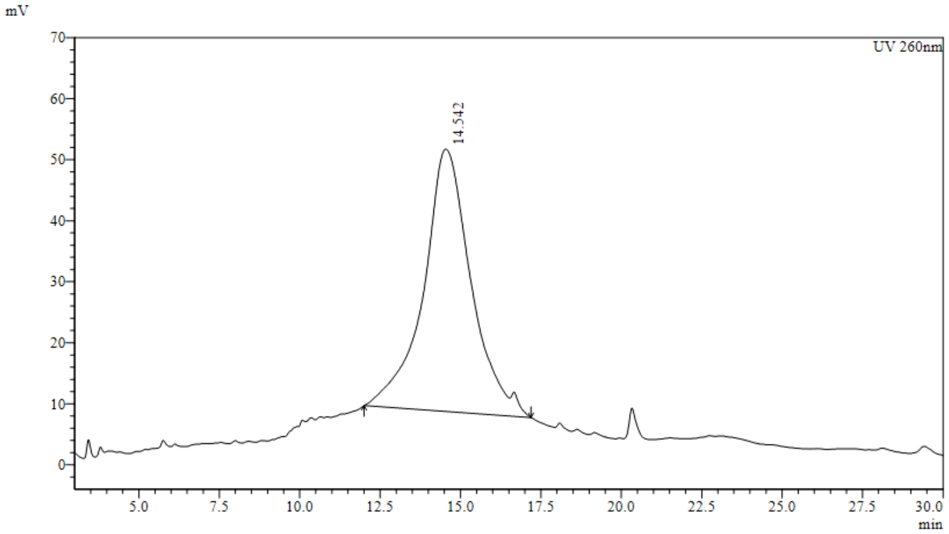


#### Peak Table

UV 260nm

| Peak# | Ret. Time | Area | Height | Area% |
|---|---|---|---|---|
| 1 | 14.542 | 4441898 | 43010 | 100.000 |
| Total | | 4441898 | 43010 | 100.000 |

**Figure S18.** HPLC chromatogram of PMO2 (after irradiation) (5′-T^A^T^A^AATTGTAACTGAGGTAAGAGG-3′)

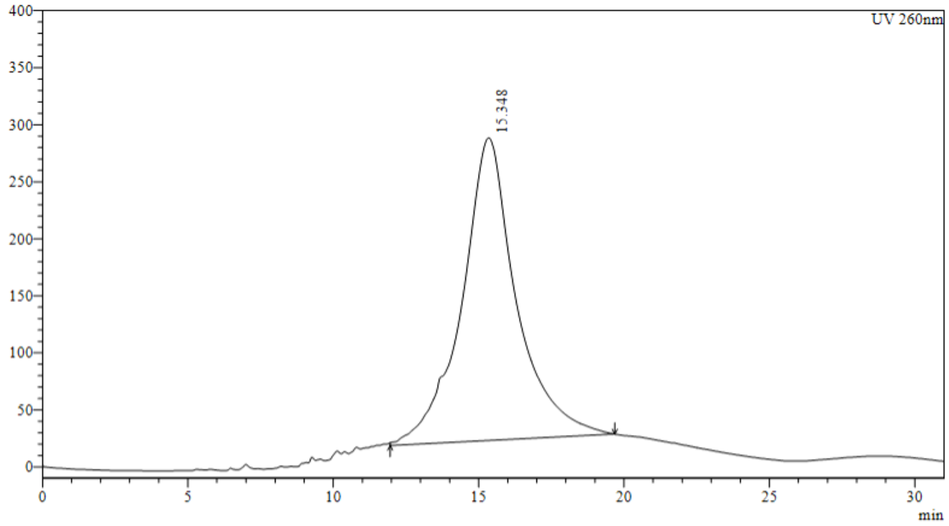


#### Peak Table

UV 260nm

| Peak# | Ret. Time | Area | Height | Area% |
|---|---|---|---|---|
| 1 | 15.348 | 34557364 | 265317 | 100.000 |
| Total | | 34557364 | 265317 | 100.000 |

**Figure S19.** HPLC chromatogram of PMO3 (5′-T^A^T^A^AATTGTAACTGAGGTAAGAGG-3′)

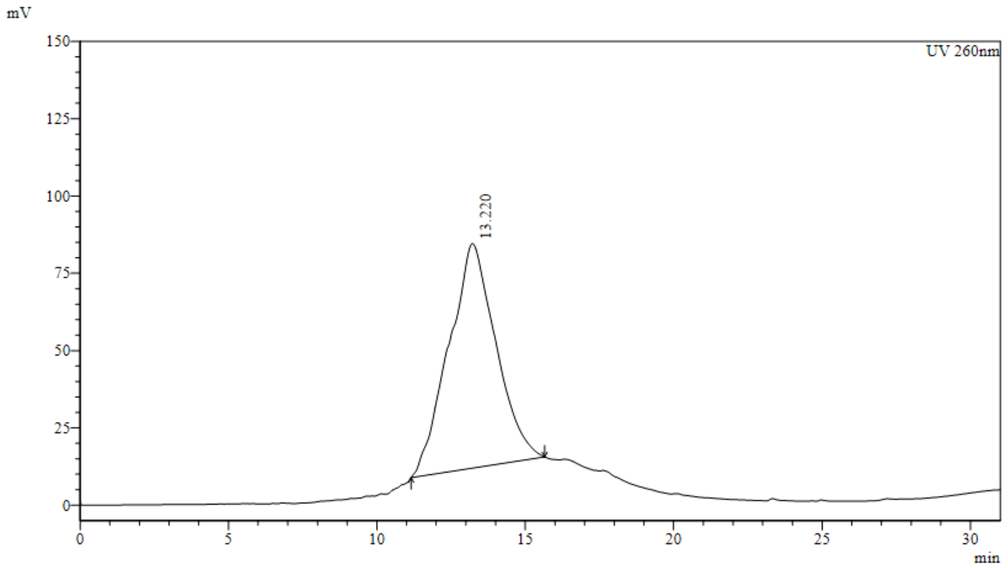


**Peak Table**

UV 260nm

| Peak# | Ret. Time | Area | Height | Area% |
|---|---|---|---|---|
| 1 | 13.220 | 8035723 | 72625 | 100.000 |
| Total | | 8035723 | 72625 | 100.000 |

**Figure S20.** HPLC chromatogram of complimentary tbMO (5'-CCTCTTACCTCAGTTACAATTTATA-3')

**Scheme S3: Synthesis of BODIPY-conjugated PMOs.**

The reported synthesized PMOs (20 nmol) were conjugated with NHS-ester of BODIPY at 3′ end in 100 µL 0.1 M aqueous $NaHCO_3$ solution containing 25% $C_2H_5OH$. The resultant mixture was vortexed for 48 h and the crude mixture was washed with DCM to remove the excess BODIPY. The resultant aqueous layer was centrifuged 3000 MWCO (Amicon ultra, Merck Milipore). Finally the purity was checked by HPLC ($\lambda_{max}$ = 498 nm).

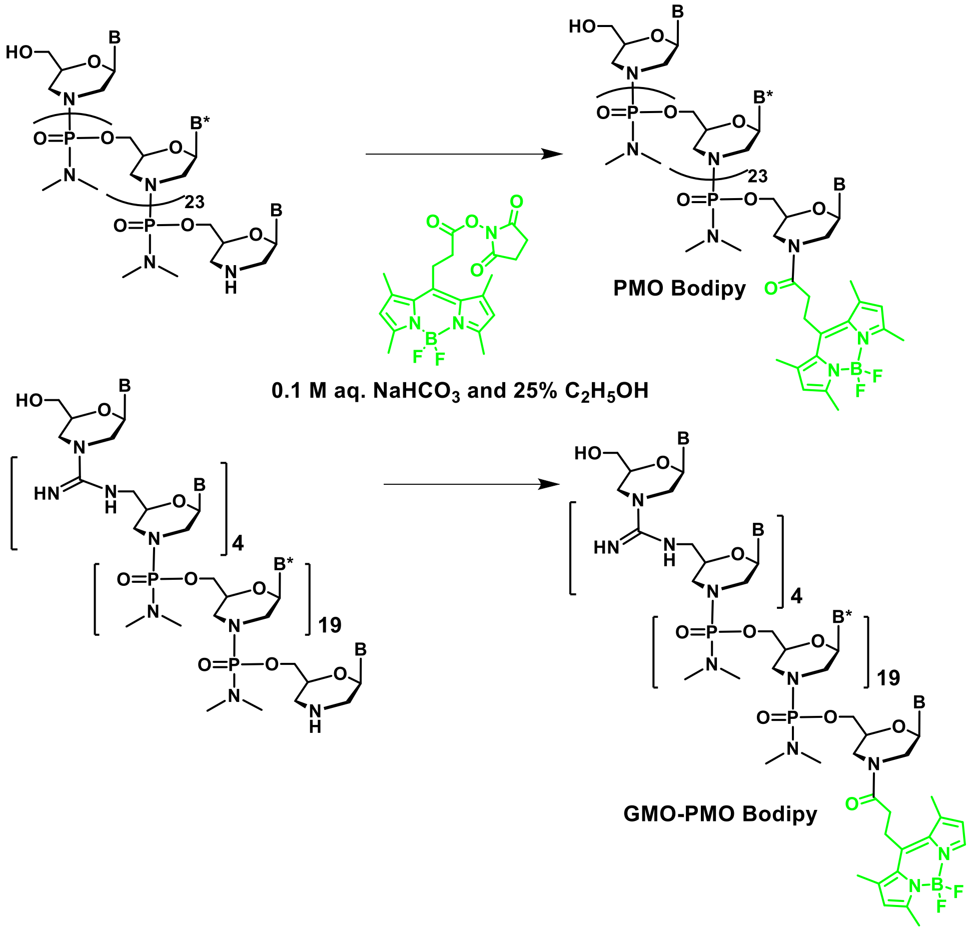

#### Chromatogram

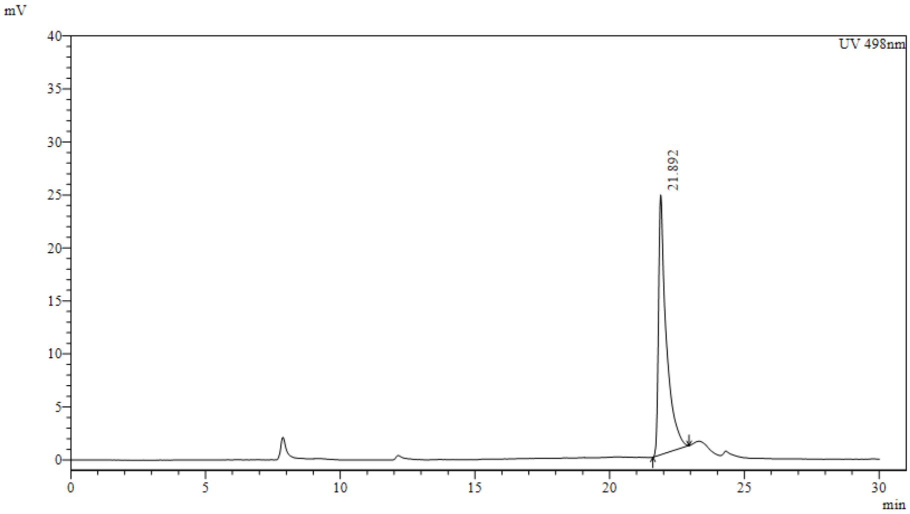


#### Peak Table

UV 498nm

| Peak# | Ret. Time | Area | Height | Area% |
|---|---|---|---|---|
| 1 | 21.892 | 499740 | 24519 | 100.000 |
| Total | | 499740 | 24519 | 100.000 |

**Figure S21.** HPLC chromatogram of PMO1-BODIPY

#### Chromatogram

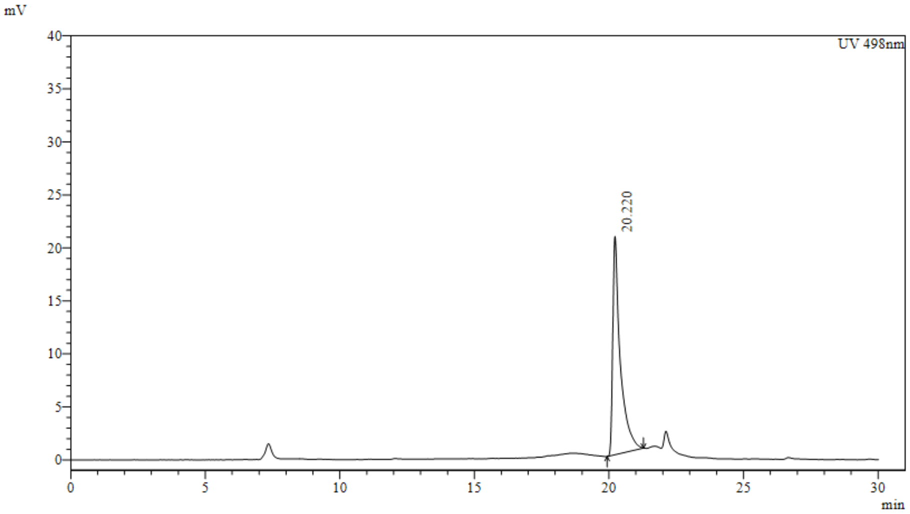


#### Peak Table

UV 498nm

| Peak# | Ret. Time | Area | Height | Area% |
|---|---|---|---|---|
| 1 | 20.220 | 396219 | 20604 | 100.000 |
| Total | | 396219 | 20604 | 100.000 |

**Figure S22.** HPLC chromatogram of PMO3-BODIPY

**Chromatogram**

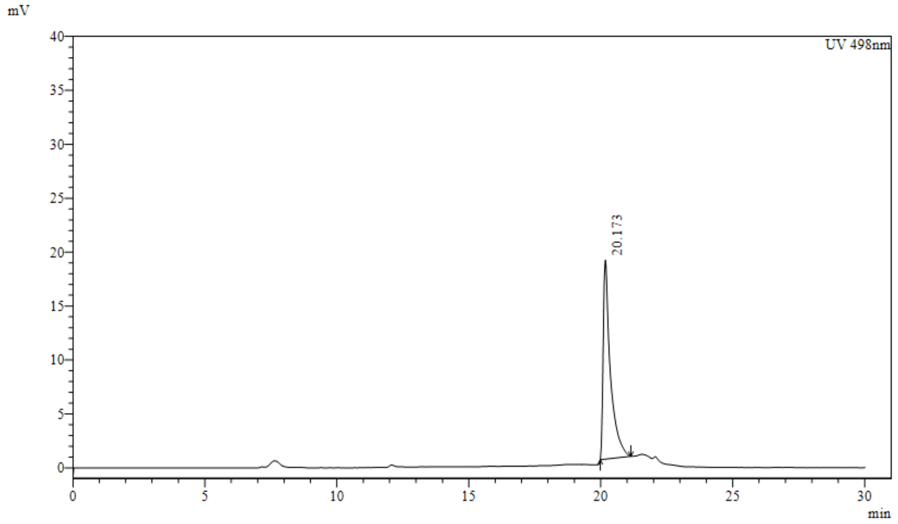


**Peak Table**

UV 498nm

| Peak# | Ret. Time | Area | Height | Area% |
|---|---|---|---|---|
| 1 | 20.173 | 343462 | 18464 | 100.000 |
| Total | | 343462 | 18464 | 100.000 |

**Figure S23.** HPLC chromatogram of PMO2-BODIPY

#### MALDI-TOF Spectra of PMOs

**Table S4**: MALDI-TOF mass of PMOs

| Sequence | Molecular formula | Calculated mass | Obtained mass |
|---|---|---|---|
| 5′-T^T^T^T^TTTTTT-3′ (GMOPMO-1)* | $C_{114}H_{170}N_{43}O_{41}P_5$ | 2952.1 | 2951.9 |
| 5′-T^A^T^A^ATTTTT-3′ (GMOPMO-2)* | $C_{114}H_{167}N_{52}O_{35}P_5KNa_3$ | 3087.1 | 3085.8 |
| 5'-TATAAATTGTAACTGAGGTAAGAGG-3' (cPMO1) | $C_{338}H_{496}N_{154}O_{111}P_{24}Na_2$ | 9281.8 | 9283.7 |
| 5′ -TATAAATTGTAACTGAGGTAAGAGG-3′ (PMO 1) | $C_{305}H_{457}N_{151}O_{96}P_{24}K_3$ | 8635.5 | 8637.1 |
| 5'-T^A^T^A^AATTGTAACTGAGGTAAGAGG-3' (cPMO2) | $C_{334}H_{480}N_{158}O_{103}P_{20}K_3$ | 9088.1 | 9088.7 |
| 5′-T^A^T^A^AATTGTAACTGAGGTAAGAGG-3′ (PMO2) | $C_{301}H_{441}N_{155}O_{88}P_{20}KNa_2$ | 8343.3 | 8341.0 |
| 5′-T^A^T^A^AATTGTAACTGAGGTAAGAGG-3′ (PMO3) | $C_{293}H_{437}N_{155}O_{88}P_{20}KNa$ | 8220.2 | 8220.1 |
| 5'-CCTCTTACCTCAGTTACAATTTATA-3'(tbMO) | $C_{291}H_{457}N_{155}O_{102}P_{24}K$ | 8186.8 | 8186.3 |

***GMO-PMO mass was detected without Trityl, accordingly their molecular formula and masses were incorporated in the table.**

a

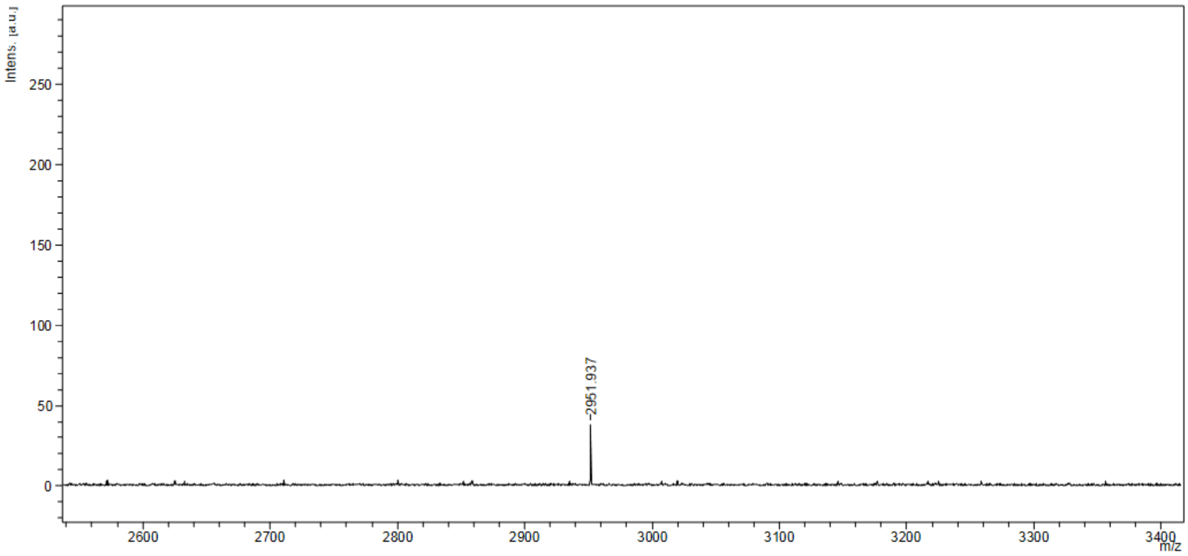


b

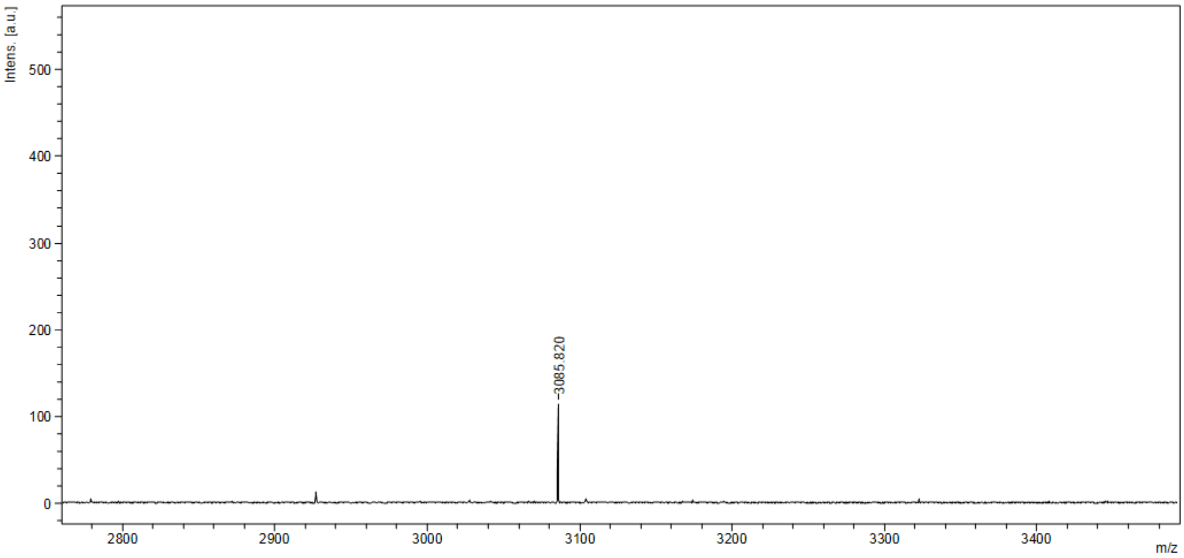


c

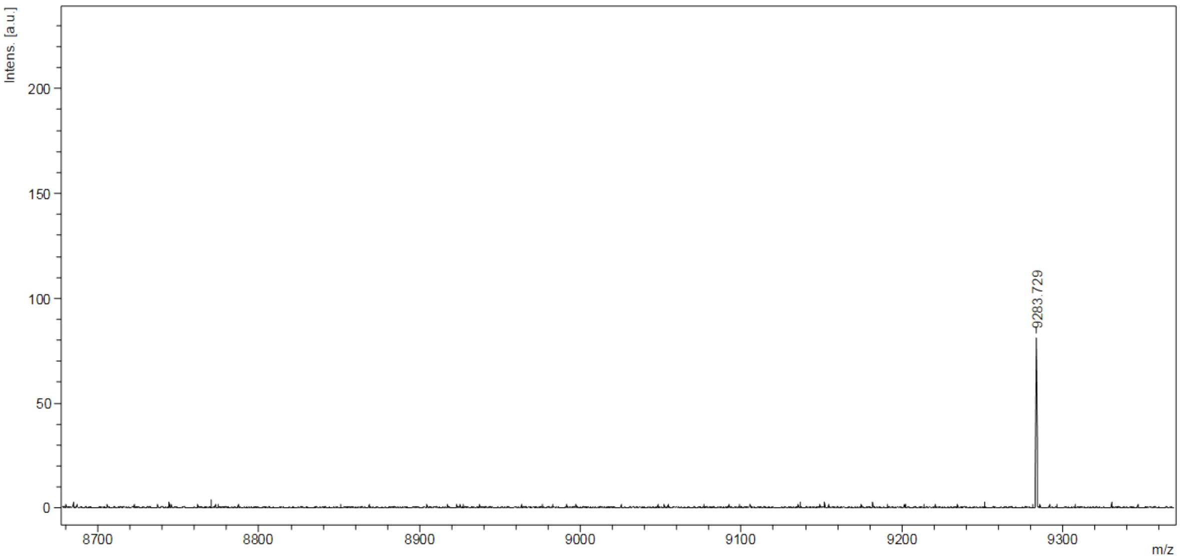


**Figure S24.** MALDI-TOF Spectra of (**a**) GMOPMO-1 (**b**) GMOPMO-2 (**c**) cPMO1

**a**

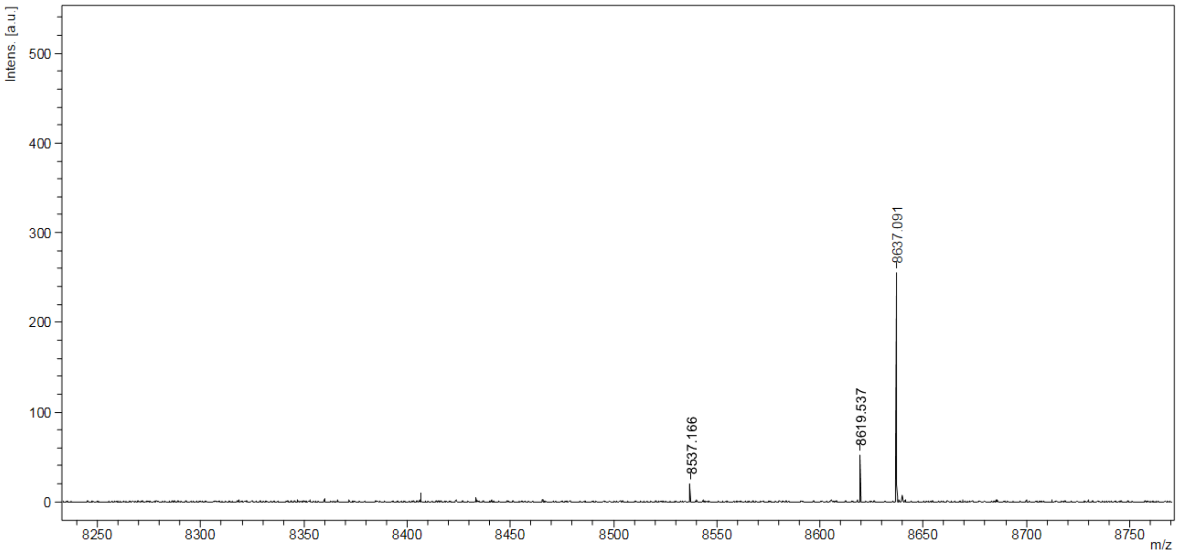


**b**

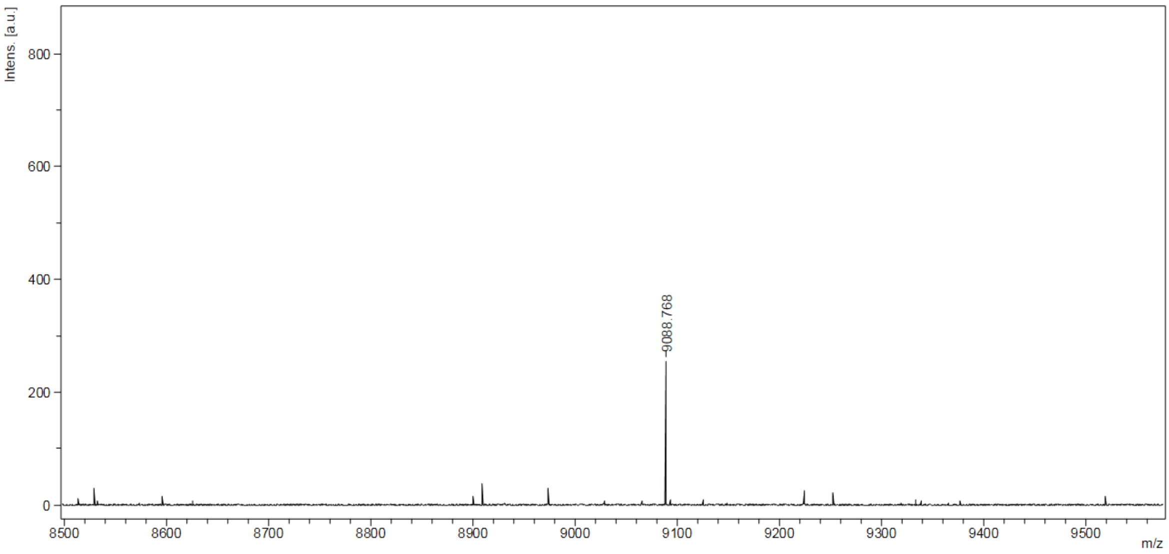


**c**

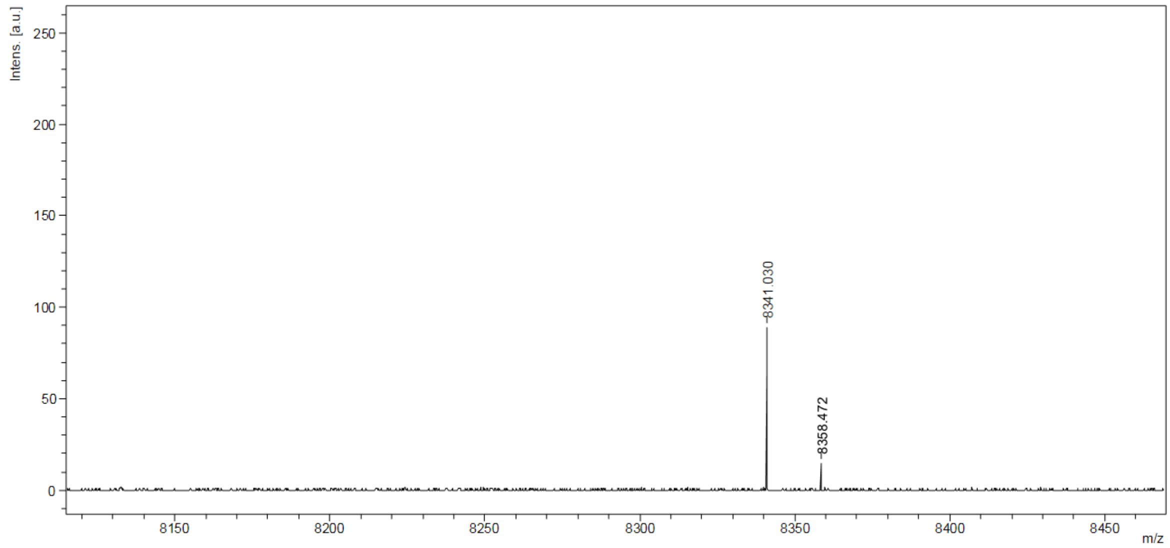


**Figure S25.** MALDI-TOF Spectra of (**a**) PMO1 (**b**) cPMO2 (**c**) PMO2

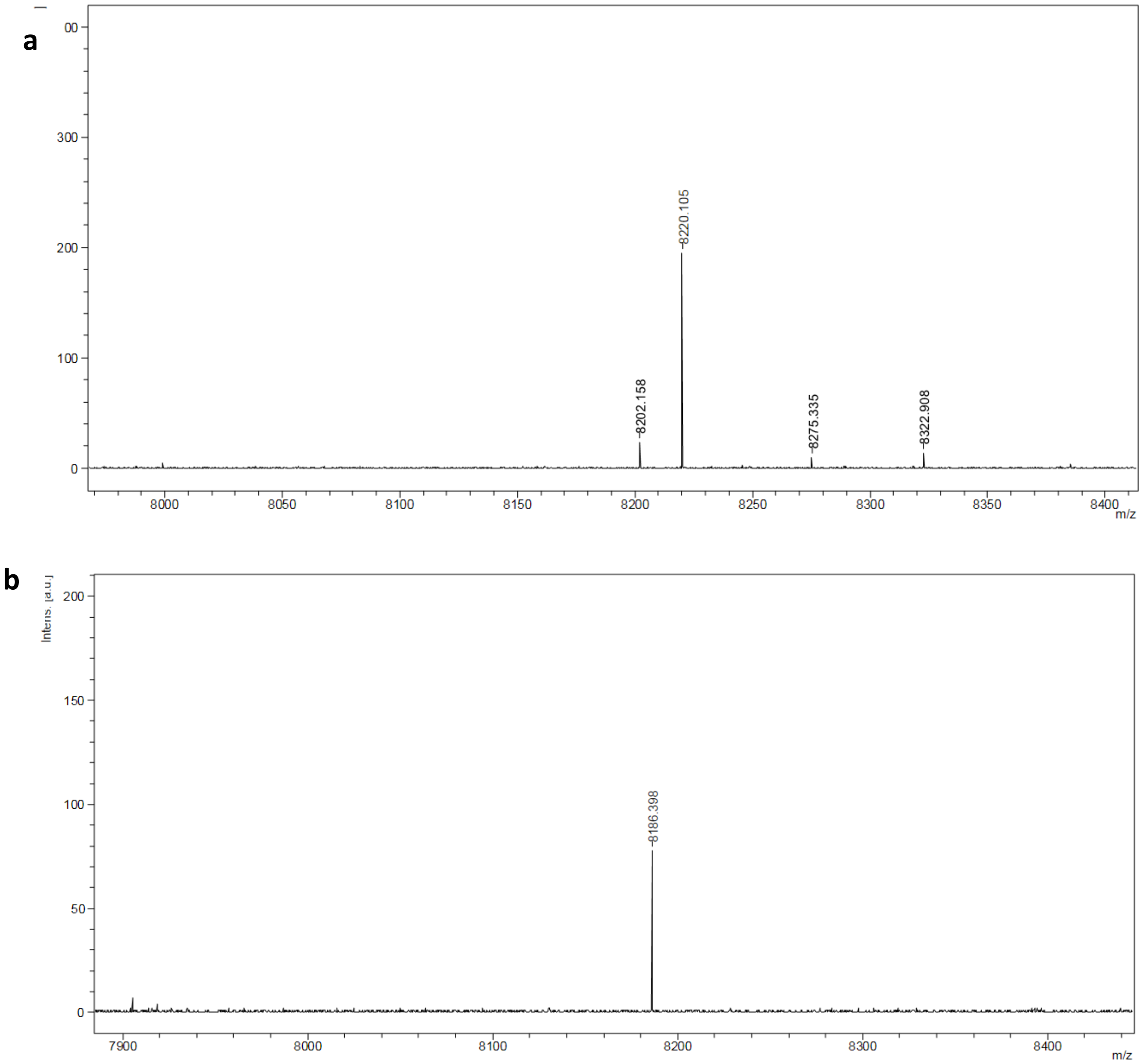


**Figure S26.** MALDI-TOF Spectra of (**a**) PMO3 (**b**) tbMO
